# Regional choroid plexus calcifications and their associations with aging, brain structure, and disease

**DOI:** 10.64898/2026.09.10.750525

**Authors:** Iyad Ba Gari, Alyssa Zhu, Ravi R Bhatt, Talia M. Nir, Neda Jahanshad

## Abstract

The choroid plexus (CP) maintains brain homeostasis through cerebrospinal fluid production and formation of the blood-cerebrospinal fluid barrier, and its dysfunction has been linked with aging and neurological disease. CP dysfunction may involve both enlargement and calcific tissue change, which may reflect distinct processes; yet population neuroimaging has focused on lateral ventricle CP volume, which cannot directly identify calcified tissue. Using quantitative susceptibility mapping (QSM) in 30,012 UK Biobank participants, we conducted, to our knowledge, the first large scale multi-ventricular study of CP calcification (CPcal), quantifying the lateral (LV), third (3rdV) and fourth (4thV) ventricles. We examined associations with age and sex, endocrine, nutritional and metabolic, mental and behavioral, nervous system, and circulatory diagnoses; family history of Alzheimer’s disease and related dementias (ADRD); cardiometabolic traits; and brain macrostructure, white matter microstructure, and subcortical susceptibility. CPcal increased with age (largest association: LV, r = 0.191) and was greater in males in the LV and 3rdV (d = 0.249-0.359) and more prevalent in the 4thV (OR = 1.274). Greater CPcal was associated with endocrine and metabolic, psychiatric, nervous system, and circulatory disorders, with the strongest and most consistent associations observed in the 3rdV (d = 0.097 - 0.125) and persisted after adjustment for ventricular volume, particularly for diabetes (d = 0.303) and tobacco use disorder (d = 0.314). Maternal ADRD family history was associated with greater LV CPcal (d = 0.063). CPcal was also associated with cortical and subcortical volumes (r = 0.033 - 0.046), white matter microstructure (r = 0.017 - 0.042), and subcortical susceptibility (r = −0.064 - 0.131). QSM-derived CPcal is a scalable, radiation-free imaging phenotype that complements volumetry by characterizing calcific CP tissue, which is associated with cardiometabolic health, brain aging, and neurodegenerative risk, particularly in the third ventricle.

## 1 Introduction

The choroid plexus (CP) is a highly vascularized epithelial tissue within the cerebral ventricles that produces the majority of cerebrospinal fluid (CSF), forms a blood-CSF barrier, and supports CSF circulation, metabolite exchange, and immune signaling [Čarna et al., 2023; Lun et al., 2015; Marques et al., 2017]. Glymphatic clearance is tightly coupled to the sleep-wake cycle and is most active during sleep, linking CSF dynamics and glymphatic function to disturbances of sleep and circadian rhythm [Christensen et al., 2022]. These functions establish the CP as an important interface between the systemic circulation and the brain and motivate its study in brain health. Structural alterations, most commonly assessed as changes in CP volume, have been reported in psychiatric and neurodegenerative disorders. CP enlargement has been observed in bipolar disorder with psychotic features and schizophrenia [Bitanihirwe et al., 2022; Lizano et al., 2019], and, in first-episode schizophrenia, larger CP volume has been associated with metabolic, neuroendocrine, and cardiovascular markers [Bitanihirwe et al., 2022; Zhou et al., 2020]. CP enlargement has also been reported across the Alzheimer’s disease spectrum [Novakova Martinkova et al., 2023]. More recently, a prospective UK Biobank study found that greater CP volume and lower CP signal intensity were associated with cognitive decline and incident dementia[Yu et al., 2026]. CP volume has been studied alongside macrostructural and microstructural brain measures in cognitively healthy adults [Alisch et al., 2021] and people with schizophrenia spectrum disorders [Yakimov et al., 2024].

However, volume alone provides only a partial characterization of CP structure. CP calcification is the most common form of physiological intracranial calcification after midlife and is typically localized to the atria of the lateral ventricles, and increases in prevalence and extent with age [Saade et al., 2019; Whitehead et al., 2015; Yalcin et al., 2016]. Although calcification has traditionally been considered an age-related finding, it may also reflect pathological processes, including dysregulated mineral metabolism, prior infection, and ventricular inflammation, that are not captured by CP volume [Whitehead et al., 2015]. Because these processes directly affect the inflammatory and secretory functions of the CP, calcification may provide a more specific marker of CP pathology than volume. Consistent with this, CPcal, but not volume, has been shown to predict cortical microglial activation, supporting its potential as a biomarker of CP pathology and neuroinflammation [Butler et al., 2023]. However, investigating CPcal in large population cohorts has remained challenging. Computed tomography (CT) exposes individuals to ionizing radiation [Sedghizadeh et al., 2012], while T1-weighted MRI cannot reliably detect calcification [Wu et al., 2009]. Consequently, CPcal has remained poorly studied in large population neuroimaging studies.

Quantitative susceptibility mapping (QSM) is a reliable MRI approach for identifying and quantifying calcification [Ruetten et al., 2019]. QSM maps are reconstructed noninvasively, providing a radiation-free imaging modality suitable for large-scale neuroimaging studies. In the UK Biobank, Wang et al. showed that voxelwise magnetic susceptibility provided information beyond conventional T2*-based measures and yielded reproducible regional susceptibility patterns across the brain [Wang et al., 2022]. To our knowledge, QSM has not yet been used to quantify CP calcification across a large population cohort or to test whether CPcal captures disease and brain-related variation beyond CP volume. Establishing these relationships would determine whether CPcal contributes complementary information to commonly used neuroimaging phenotypes.

Here, we address this gap by studying 30,012 middle-aged and older adult participants from the UK Biobank [Bycroft et al., 2018; Miller et al., 2016]. As the CP differs in cellular composition and physiological function across the ventricles, CPcal in each region may reflect distinct underlying pathophysiology [Dani et al., 2021]. Therefore, we quantified calcification separately in the lateral, third, and fourth ventricles. In our study, we (i) characterized age and sex effects on CPcal across the three ventricular regions; (ii) tested whether calcification is associated with aging-related metabolic, neurological, psychiatric, and cerebrovascular disorders previously linked to CP volume; (iii) evaluated associations with cardiometabolic risk factors and risk factors for Alzheimer’s disease and related dementias (ADRD); and (iv) examined associations between CPcal and regional brain macrostructure, white matter microstructure and iron deposition.

## 2 Methods

### 2.1 UK Biobank Participants Information

We analyzed multimodal neuroimaging data from 30,012 participants in the UK Biobank dataset (downloaded in 2021) [Miller et al., 2016] as part of a cross-sectional analysis. Among 500,000 UK Biobank participants, 38,459 had imaging data available; after excluding 3,354 with the flipped SWI acquisition protocol and 5,093 with severe Gibbs artifacts in QSM images, 30,012 were included in the final analysis. All the individuals included in this study had full brain T1-weighted, diffusion-weighted, and susceptibility-weighted images (T1w, DWI, and SWI respectively).

### 2.2 Imaging Acquisition and Processing

All T1-weighted, susceptibility-weighted, and diffusion-weighted MRIs were acquired by the UK Biobank according to the protocol described by Miller et al. (2016)[Miller et al., 2016]. A summary of the image acquisition procedures and subsequent processing steps is provided below. Diffusion fractional anisotropy (FA) and mean diffusivity (MD) measures were obtained from the UK Biobank image-derived phenotypes (IDPs). FreeSurfer-derived cortical and subcortical metrics, quantitative susceptibility maps (QSM), and CPcal volumes were derived using the image-processing pipeline described below.

#### 2.2.1 T1-weighted MRI

T1-weighted brain images (T1w) were acquired using a 3D MPRAGE sequence with the following parameters: voxel size = 1×1×1 mm^3^, field of view = 208×256×256, in-plane acceleration iPAT=2, sagittal, total scan time = 5 minutes. We converted T1w DICOMs to NIfTI and then processed them using FreeSurfer 7.1’s recon_all pipeline [Fischl, 2012]. Regional cortical surface area and cortical thickness were extracted for each hemisphere and averaged across the left and right hemispheres for analysis. Volumes were also extracted for seven bilateral subcortical regions of interest (ROIs): thalamus, caudate, putamen, pallidum, hippocampus, amygdala, and accumbens. We also extracted the masks for the lateral ventricles (LV), 3rd ventricle (3rdV), and 4th ventricle (4thV) from the Freesurfer outputs to use as a mask for localized calcification estimations. The T1w image and corresponding FreeSurfer labels were linearly registered to the SWI magnitude image using FSL’s flirt with 12 degrees of freedom (dof). The SWI magnitude image is in the same space as the QSM image and provides anatomical contrast for reliable registration.

#### 2.2.2 Quantitative susceptibility mapping and defining the choroid plexus calcification

Quantitative susceptibility maps (QSM) were derived from DICOMs of the susceptibility-weighted images (SWI). SWI scans were collected on 3T Siemens Skyra scanners using a standard Siemens 32-channel RF receive head coil (magnitude and phase images were saved for each RF coil and echo time separately). SWI MRI data were acquired using a double-echo GRE sequence with echo times (TE) = 9.42/20 ms, voxel size = 0.8×0.8×3 mm^3^, in-plane acceleration iPAT=2, axial, total scan time = 2.5 minutes. Further details about UK Biobank imaging protocols are described in Miller et al 2016 [Miller et al., 2016].

QSM images were reconstructed using the UK Biobank QSM pipeline (https://git.fmrib.ox.ac.uk/cwang/uk_biobank_qsm_pipeline), developed by Wang et al. (2022) [Wang et al., 2022]. First, phase images from the 32 channels were combined using the MCPC-3D-S approach [Eckstein et al., 2018]. The combined phase images were then unwrapped using a Laplacian-based algorithm implemented in the STI Suite toolbox [Schofield and Zhu, 2003]. Brain masks, generated from the standard UK Biobank pipeline [Du et al., 2018], were applied to exclude voxels with unreliable phase values. A local field map was subsequently obtained by removing background field contributions using the variable-kernel sophisticated harmonic artifact reduction for phase data (v-SHARP) method with a kernel size of 12 mm [Schweser et al., 2011]. Finally, dipole inversion was performed using the iLSQR algorithm [Li et al., 2015], also provided by the STI Suite toolbox.

QSM images underwent visual quality control for Gibbs ringing, motion artifacts, and visible pathology. Artifacts were graded as absent (0), mild (1), or moderate to severe (2). All grade-2 images and images with pathology that could compromise the analysis were excluded, whereas grades 0-1 were retained. When present, Gibbs ringing was distributed throughout the image rather than localized to a specific anatomical region. Each CP region was therefore inspected directly, and images were excluded if artifacts obscured the region of interest. In retained images, mild ringing did not prevent CPcal assessment, although residual measurement error in subcortical susceptibility estimates cannot be excluded.

Physiological calcification of the CP was extracted by intensity thresholding of the QSM images that passed quality control based on the negative correlation between CT attenuation and QSM reported in the CP by Oshima et al. (2020) [Oshima et al., 2020] and visual inspection QSM images. Calcification was quantified within the lateral (LV), third (3rdV), and fourth (4thV) ventricle masks derived from FreeSurfer (Section 2.2.1) after registration to the QSM space. Within each ventricle mask, voxels with magnetic susceptibility below −0.1 ppm (−100 ppb) were labeled as calcification. CPcal volume was computed from the number of voxels and the QSM voxel volume and is reported in mm^3^.

The distribution of CPcal volumes in the lateral ventricles and third ventricle volumes were right-skewed (Figure 1A), so, for all analyses, we applied a log transformation for these measures. CPcal in the fourth ventricle was analyzed as a binary metric, categorized as calcified or not, due to 66% of participants having no calcification in the fourth ventricle (Table 1).

**Figure 1.**
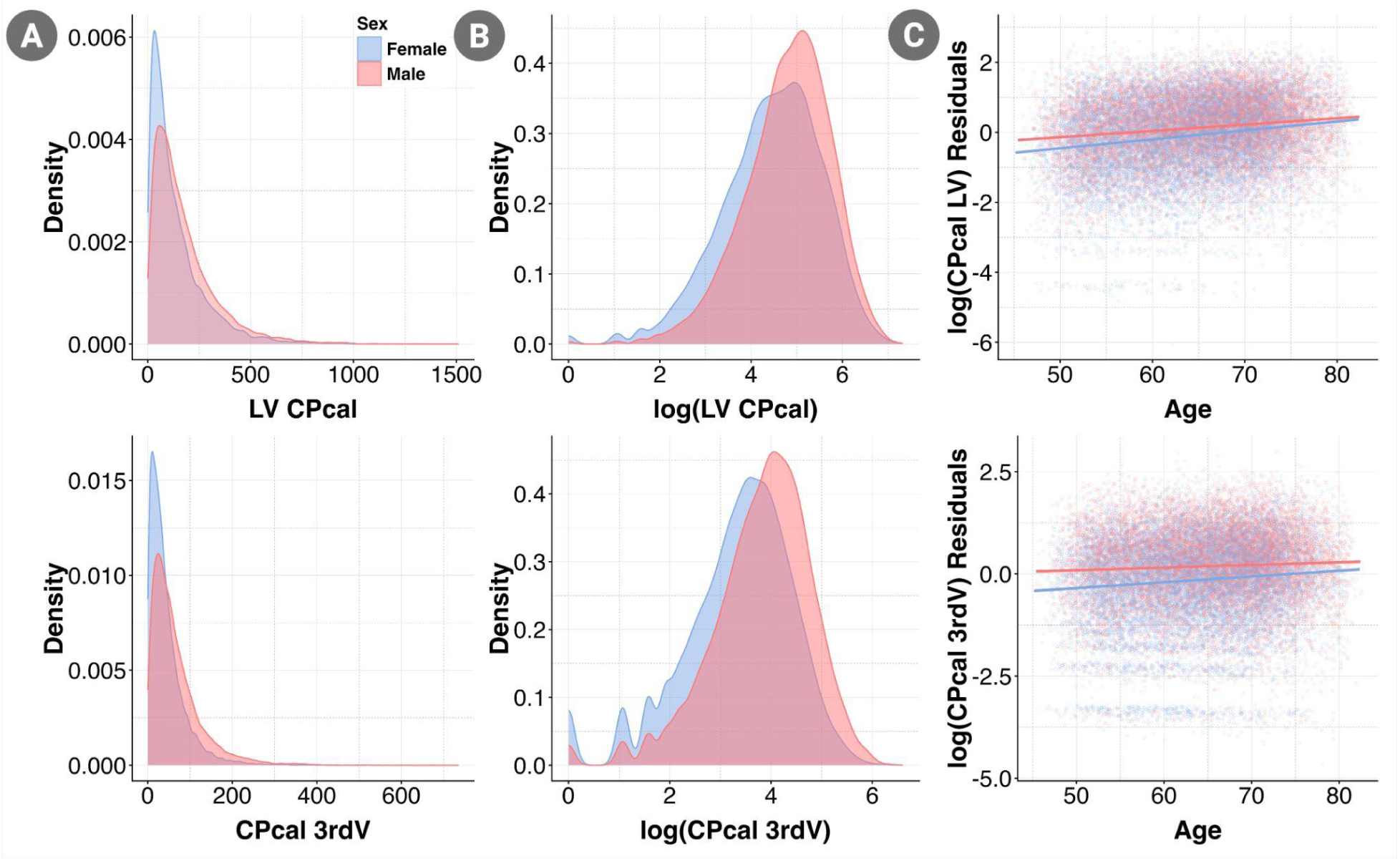
**A.** Distribution of CPcal volumes in lateral (LV) and third ventricles (3rdV) stratified by sex. **B.** Log-transformed CPcal volumes. **C.** Scatterplot illustrating the association of log-transformed CPcal volumes with age and stratified by sex, after adjusting for lateral ventricle volume, intracranial volume (ICV), scanner table position, and imaging site.

**Table 1.**
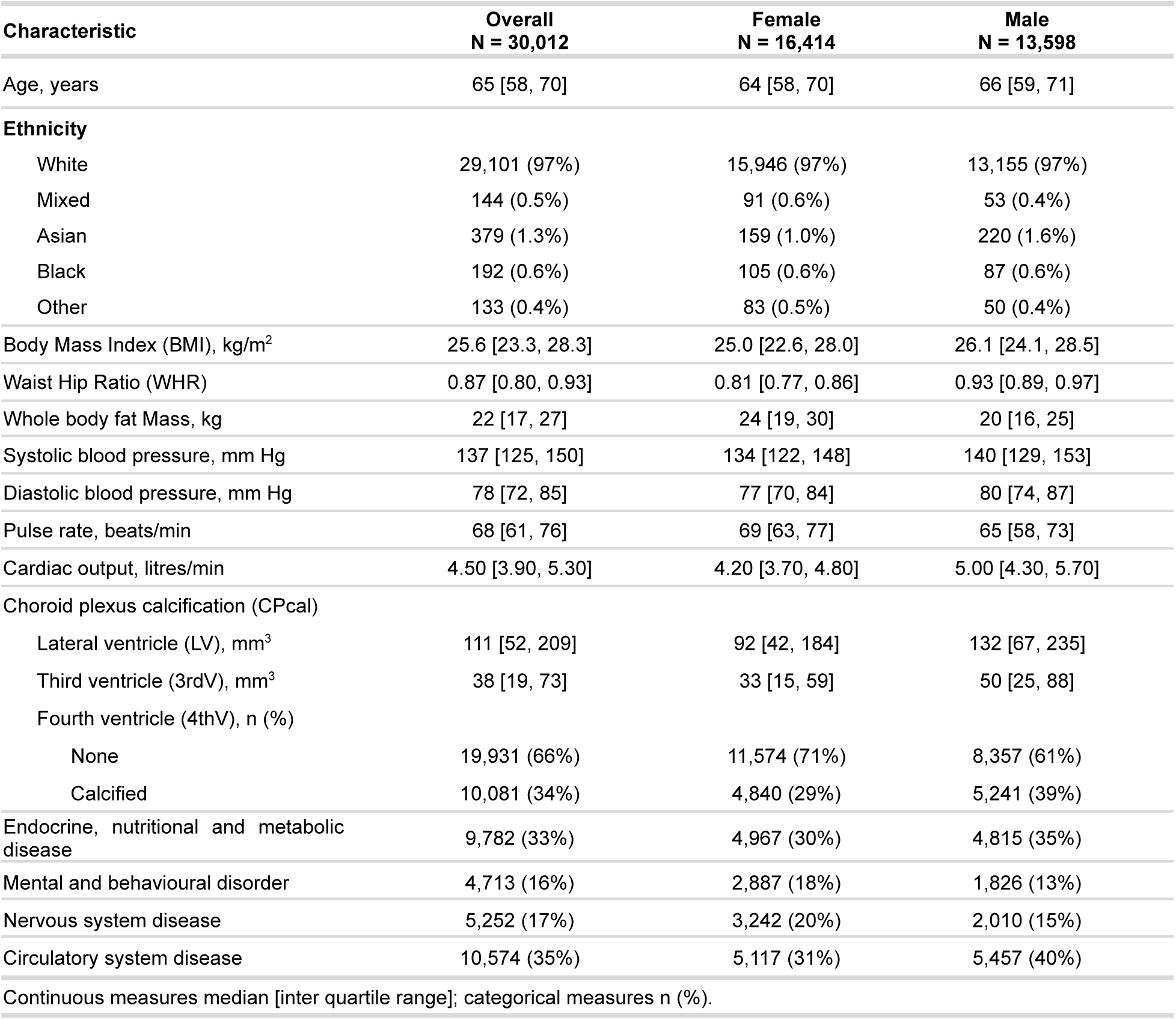
Demographic, cardiometabolic, and disease characteristics of the UK Biobank study cohort (N = 30,012), overall and stratified by sex. Continuous variables are median [interquartile range (IQR)] and categorical variables are n (%). CPcal volumes are reported in mm^3^.

#### 2.2.3 Diffusion-weighted MRI

dMRI scans were acquired with 2×2×2 mm^3^ voxels and included 50 b=1000 s/mm^2^ and 50 b=2000 s/mm^2^ diffusion-weighted images, and 5 b0 volumes. The processing has been described by Alfaro-Almagro and colleagues [Alfaro-Almagro et al., 2018]. We used UK Biobank-provided image-derived phenotypes for mean fractional anisotropy (FA) and mean diffusivity (MD) within white matter tracts defined on the skeleton using the Johns Hopkins University white matter atlas (JHU).

### 2.3 Statistical Analyses

#### 2.3.1 Age and sex effects on log-transformed CPcal volumes

Associations between log-transformed CPcal volume in the LV and 3rdV (dependent variable) and age were tested with linear regressions. Associations between CPcal presence in the 4thV (i.e. calcified vs not) was evaluated with logistic regression. All models were adjusted for sex, lateral ventricle volume, intracranial volume (ICV), scanner table position, and imaging site; this covariate set was used in all subsequent analyses unless otherwise specified. These confounds are selected based on previous research [Alfaro-Almagro et al., 2021]. We additionally analyzed the age-by-sex interaction and repeated the age analysis stratified by sex to evaluate CPcal differences between males and females. Multiple comparisons were controlled with the Benjamini-Hochberg (BH) false discovery rate (FDR, q = 0.05) across the three CPcal regions. For CPcal in the lateral and third ventricles, we also performed quantile regression at the 5th, 25th, 50th, 75th, and 95th percentiles of the calcification distribution to assess whether associations varied across the outcome distribution. Quantile-regression results are reported in Supplementary Figures S1-S2 and Supplementary Tables S06-S10.

#### 2.3.2 Disease and risk factor effects on log-transformed CPcal volumes

We compared CPcal across participants with and without diagnoses in four ICD-10 chapters: endocrine, nutritional and metabolic diseases (chapter IV), mental and behavioral disorders (V), diseases of the nervous system (VI), and diseases of the circulatory system (IX). These chapters were selected as prior studies have linked CP structural abnormalities to metabolic and cardiovascular burden [Li et al., 2024; Zhou et al., 2020], psychiatric disorders [Bitanihirwe et al., 2022; Lizano et al., 2019], and neurological or neurodegenerative disease [Choi et al., 2022; Egorova et al., 2019]. Healthy controls were participants with no recorded diagnosis in any of the four chapters (n = 11,480). For each disease comparison, a separate 1:1 matched control set was drawn from this control pool; thus, controls varied across comparisons, and individuals could be selected more than once. Diagnoses were derived from ICD-10 codes obtained from three sources:

1. Hospital in-patient records (IDs <u>41202</u>, <u>41270</u>) and their corresponding dates (IDs <u>41262</u>, <u>41280</u>). We simplified the ICD-10 codes to a three-tier system: chapter, level-1, and specific diseases, by truncating the original codes. The dates corresponding to the ICD-10 diagnoses were then compared with the dates of subjects’ visits to assessment centers (ID <u>53</u>). This comparison determined whether a diagnosis was present at the time of the imaging visit.
2. Self-reported health data (ID <u>41204</u>) and associated dates (ID <u>20008</u>). We transformed these self-reports into ICD-10 codes using UK Biobank defined data-coding ID <u>609</u>. Self-report dates, recorded as a decimal year (e.g., 2008.5 denotes mid-2008), were converted to approximate calendar dates, excluding records with years earlier than 1930, uncertain/unknown dates (−1), or when subjects preferred not to answer (−3). These dates were then compared with the dates of attendance at assessment centers (ID <u>53</u>) to assess disease presence at that time.
3. Death register data (ID <u>40001</u>) and corresponding dates of death (ID <u>40000</u>). The process for these data was similar to that of hospital in-patient records, with ICD-10 codes truncated to identify chapter, level-1, and specific diseases. These dates were then compared with the subjects’ assessment center visit dates (ID <u>53</u>) to determine disease presence.

Further details of the level-1 and disorder classifications are provided in Table S1. For each disease contrast, cases were matched 1:1 to healthy controls on age and sex using nearest-neighbour propensity-score matching (logistic-regression distance; MatchIt) [Ho et al., 2011]. In a progression analysis, cases were restricted to participants first diagnosed after their baseline imaging visit and were compared with matched controls who remained free of these diagnoses throughout follow-up. To verify that the associations with log-transformed third ventricle CPcal volume were not an artifact from ventricular enlargement, we re-fitted all third ventricle disease models with FreeSurfer-derived third ventricle cerebrospinal fluid volume included as an additional covariate.

Further, we explored the effect of pulse rate, cardiac output, basal metabolic rate (BMR), whole body fat mass (BFM), waist-to-hip ratio (WHR), and body mass index (BMI) on log-transformed CPcal volumes. Disease contrasts (dichotomous matched case/control) and the continuous cardiovascular and metabolic risk factors were analyzed with the covariate set described above, additionally including the age-by-sex interaction. FDR correction (q = 0.05) was applied across all tests within each analysis family - the three CPcal regions crossed with every disease or risk-factor trait examined, rather than across only three CPcal regions.

Individuals diagnosed with Alzheimer’s disease or dementia cases were sparse in this imaging cohort. Therefore, we used participant reported parental history of Alzheimer’s disease or dementia as a proxy phenotype for familial ADRD susceptibility which is consistent with prior UK Biobank studies [Liu et al., 2017; Marioni et al., 2018]. This proxy reflects familial susceptibility rather than an ADRD diagnosis in the participant and may capture shared genetic and environmental influence. Models were adjusted for age, sex, age-by-sex interaction, lateral ventricle volume, intracranial volume, scanner table position, and imaging site. To reduce ancestry-related confounding, the analysis was restricted to 26,072 participants who self-reported European ethnicity, excluding 3,940 participants who self-reported non-European ethnicity. We identified individuals with a family history of ADRD from family history in the UK Biobank dataset by examining the recorded illnesses of the father (IDs 20107) and mother (IDs 20110) using data-coding (ID <u>1010</u>) for which the code value ‘10’ indicates the presence of a family history. Participants were classified as controls when neither parent had a recorded ADRD diagnosis and the parental-illness fields indicated none of the listed conditions (data-coding 1010, values −17 and −27 for father and mother, respectively). Cases were matched 1:1 to controls on age and sex, and FDR correction (q = 0.05) was applied across the three CPcal regions and all family-history comparisons.

#### 2.3.3 Brain Volume and Microstructural Associations with CPcal Volume

To examine associations between CPcal volume and brain structure and microstructure, we regressed each regional measure - cortical thickness and surface area, subcortical volume, subcortical magnetic susceptibility (QSM), and white matter fractional anisotropy (FA) and mean diffusivity (MD) - on log-transformed CPcal volume (or, for the 4th ventricle, calcified vs not), adjusting for the covariates described above. False discovery rate (FDR) correction (q = 0.05) was applied within each measure, separately for cortical thickness, surface area, subcortical volume, FA, and MD.

#### 2.3.4 Computing the effect sizes (Cohen’s d, partial r, and odds ratios)

Effect sizes were computed from t-values obtained in multiple linear regression models. For dichotomous predictors, t-values were converted to Cohen’s d using the following formula:

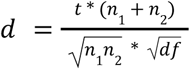

Where *n*_1_, and *n*_2_ are the sample sizes of the two groups, and *df* is the residual degrees of freedom.

For continuous predictors, effect sizes were estimated by converting t-values to partial correlation coefficients (*r*):

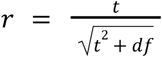

In both cases, *df* represents the residual degrees of freedom from the regression model.

Odds ratios (ORs) were reported as standardized effect sizes in logistic regression models, derived from the model coefficients as:

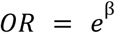

where β is the estimated regression coefficient.

## 3 Results

Our study cohort included 30,012 participants (16,414 female and 13,598 male) and summarized in Table 1.

### 3.1 Choroid plexus calcification volume associations with age and sex

Across participants, CPcal volumes in all three ventricular regions were positively associated with age, after adjusting for sex, lateral ventricle volume, intracranial volume (ICV), scanner table position, and imaging site. The largest regional effect was observed in the LV (r = 0.191, p = 1.92×10^−236^). On average, males had larger CPcal volumes than females in the lateral and third ventricles (d= 0.248, p = 1.2×10^−97^, and d = 0.358, p = 1.5×10^−199^ respectively). A significant age-by-sex interaction was observed for CPcal volume in the LV (r = −0.0208) and 3rdV (r = −0.0225), but not the 4thV (Table 2). CPcal was only modestly correlated with FreeSurfer-derived choroid plexus volume (Pearson r = 0.221, p = 3.3×10^−329^ for the LV; r = 0.240, p = 6.7×10^−388^ for the 3rdV).

**Table 2.** Associations between choroid plexus calcification (CPcal) volumes and demographic variables (age, sex, and age-by-sex interaction) in 30,012 participants. Linear regression models were used for CPcal in the lateral ventricles (LV) and third ventricle (3rdV), and logistic regression for the fourth ventricle (4thV) due to its binary distribution (presence vs. absence of calcification).

| Demographic Variable | CPcal region | Effect Size | $\beta$ | SE | 95% CI | P-value |
| --- | --- | --- | --- | --- | --- | --- |
| Age | LV | 0.1910 | 0.0286 | $8.62 \times 10^{-4}$ | (0.0269, 0.0302) | **$1.92 \times 10^{-236}$** ** |
| Sex |  | 0.2485 | 0.2745 | 0.0130 | (0.2489, 0.3000) | **$1.16 \times 10^{-97}$** ** |
| Age-by-Sex | | -0.0208 | $-5.49 \times 10^{-3}$ | 0.0015 | (-0.0085, -0.0025) | **$3.96 \times 10^{-4}$** ** |
| Age | 3rdV | 0.0971 | 0.0146 | $8.79 \times 10^{-4}$ | (0.0129, 0.0163) | **$1.05 \times 10^{-61}$** ** |
| Sex |  | 0.3585 | 0.4041 | 0.0133 | (0.3780, 0.4302) | **$1.53 \times 10^{-199}$** ** |
| Age-by-Sex | | -0.0225 | $-6.06 \times 10^{-3}$ | $1.58 \times 10^{-3}$ | (-0.0092, -0.0030) | **$1.26 \times 10^{-4}$** ** |
| Age | 4thV | 1.0112 | 0.0111 | $1.84 \times 10^{-3}$ | (1.0075, 1.0149) | **$1.63 \times 10^{-9}$** ** |
| Sex |  | 1.2736 | 0.2418 | 0.0277 | (1.2063, 1.3446) | **$2.4 \times 10^{-18}$** ** |
| Age-by-Sex | | 1.0064 | $6.34 \times 10^{-3}$ | $3.32 \times 10^{-3}$ | (0.9998, 1.0129) | 0.0562 |

\*\*Uncorrected $p \leq$ critical FDR threshold \*Uncorrected $p \leq 0.05$

### 3.2 Disease and Risk Factor Associations with CPcal

Compared with age- and sex-matched healthy controls, CPcal volumes were significantly larger in participants diagnosed with endocrine, nutritional and metabolic diseases (ICD-10 chapter IV), mental and behavioral disorders (chapter V), diseases of the nervous system (chapter VI), and diseases of the circulatory system (chapter IX) (Table 3). Endocrine and metabolic disease showed greater CPcal volume in the LV (d = 0.0386, p = 0.0150) and the 3rdV (d = 0.1214, p = 2.23×10^−14^). Mental and behavioral disorders showed greater CPcal volume in the LV (d = 0.0582, p = 0.0053) and the 3rdV (d = 0.1247, p = 2.49×10^−9^). Nervous-system disease showed greater CPcal volume in the LV (d = 0.0447, p = 0.0242) and the 3rdV (d = 0.0975, p = 9.10×10^−7^). Circulatory-system disease showed greater CPcal volume in the 3rdV (d = 0.1128, p = 4.75×10^−13^). Effects were consistently strongest in the 3rdV, and no diagnostic chapter showed a significant difference in the 4thV (Table 3). Follow-up analyses examined level-1 (block) categories within each chapter. Quantile regression supported the primary findings, with most associations remaining significant at one or more centiles and no change in effect direction. Disease effects were largely consistent across the calcification distribution, while age and cardiometabolic effects varied more across centiles. Full results are provided in Supplementary Figures S1–S2 and Tables S06–S10.

**Table 3.** Associations between choroid plexus calcification (CPcal) volumes and ICD-10 diagnostic chapters, focusing on endocrine and metabolic disorders, mental and behavioral disorders, diseases of the nervous system, and diseases of the circulatory system. Linear regression models were used for CPcal in the lateral ventricles (LV) and third ventricle (3rdV), and logistic regression for the fourth ventricle (4thV) due to its binary distribution (calcified vs not calcified).

| ICD-10 chapters | CPcal Region | Effect Size | $\beta$ | SE | 95% CI | P-value |
| --- | --- | --- | --- | --- | --- | --- |
| Endocrine, nutritional and metabolic diseases (N=7939) | LV | 0.0386 | 0.0386 | 0.0159 | (0.0075, 0.0698) | **0.0150\*\*** |
|  | 3rdV | 0.1214 | 0.1258 | 0.0165 | (0.0935, 0.1581) | **2.23x10<sup>-14\*\*</sup>** |
|  | 4thV | 1.0270 | 0.0266 | 0.0346 | (0.9596, 1.0990) | 0.4423 |
| Mental and behavioral disorders (N=4589) | LV | 0.0582 | 0.0605 | 0.0217 | (0.0179, 0.1030) | **5.33x10<sup>-3\*\*</sup>** |
|  | 3rdV | 0.1247 | 0.1330 | 0.0223 | (0.0893, 0.1767) | **2.49x10<sup>-9\*\*</sup>** |
|  | 4thV | 1.0696 | 0.0673 | 0.0459 | (0.9776, 1.1703) | 0.1428 |
| Diseases of the nervous system (N=5086) | LV | 0.0447 | 0.0463 | 0.0205 | (0.0060, 0.0865) | **0.0242\*\*** |
|  | 3rdV | 0.0975 | 0.1027 | 0.0209 | (0.0617, 0.1436) | **9.1x10<sup>-7\*\*</sup>** |
|  | 4thV | 1.0690 | 0.0667 | 0.0433 | (0.9821, 1.1637) | 0.1232 |
| Diseases of the circulatory system (N=8248) | LV | 0.0309 | 0.0310 | 0.0156 | (4.16x10<sup>-4</sup>, 0.0615) | 0.0470 |
|  | 3rdV | 0.1128 | 0.1162 | 0.0161 | (0.0848, 0.1477) | **4.75x10<sup>-13\*\*</sup>** |
|  | 4thV | 1.0050 | 0.0050 | 0.0340 | (0.9402, 1.0742) | 0.8835 |

\*\*Uncorrected $p \leq$ critical FDR threshold \*Uncorrected $p \leq 0.05$

#### 3.2.1 Endocrine, nutritional and metabolic diseases

Within ICD-10 chapter IV, larger CPcal volumes were consistently associated with diabetes mellitus, obesity, and metabolic disorders, and were most significant in the 3rdV (Table 4). Participants with diabetes mellitus had greater CPcal in the LV (d = 0.1084, p = 0.0047) and the 3rdV (d = 0.3030, p = 3.89×10^−15^). Among diabetes subtypes, non-insulin-dependent diabetes was associated with greater CPcal in the 3rdV (d = 0.2875, p = 5.68×10^−11^); unspecified diabetes was associated with greater CPcal in the LV (d = 0.1387, p = 0.0032) and the 3rdV (d = 0.3391, p = 7.47×10^−13^); insulin-dependent diabetes was associated with greater CPcal in the 3rdV (d = 0.3270, p = 0.0104). Obesity and other hyperalimentation was associated with greater CPcal in the LV (d = 0.1219, p = 0.0114) and the 3rdV (d = 0.1743, p = 2.99×10^−4^). The specific diagnosis of obesity showed a significant association in the 3rdV (d = 0.1706, p = 4.08×10^−4^), with the LV association not surviving FDR correction. Metabolic disorders were associated with greater CPcal in the 3rdV (d = 0.1256, p = 1.21×10^−11^); the LV association did not survive FDR correction. Among metabolic disorders, disorders of lipoprotein metabolism and other lipidaemias showed the strongest associations, with greater CPcal in the LV (d = 0.0529, p = 0.0059) and the 3rdV (d = 0.1379, p = 7.13×10^−13^). Volume depletion was also associated with greater CPcal in the 3rdV (d = 0.3331, p = 0.0029). No significant associations were observed for disorders of thyroid gland, other endocrine glands, or other nutritional deficiencies in any ventricle.

**Table 4.** Associations between choroid plexus calcification (CPcal) volumes and level-1 diagnostic categories within ICD-10 chapter IV (Endocrine, nutritional and metabolic diseases). Significant level-1 categories are further broken down into specific ICD-10 disorders. Linear regression models were used for CPcal in the lateral ventricles (LV) and third ventricle (3rdV), and logistic regression for the fourth ventricle (4thV) due to its binary distribution (calcified vs not calcified).

| ICD-10 chapters | CPcal Region | Effect Size | $\beta$ | SE | 95% CI | P-value |
| --- | --- | --- | --- | --- | --- | --- |
| Disorders of thyroid gland (N=2350) | LV | 0.0200 | 0.0205 | 0.0299 | (-0.0381, 0.0792) | 0.4928 |
|  | 3rdV | 0.0463 | 0.0488 | 0.0308 | (-0.0115, 0.1091) | 0.1128 |
|  | 4thV | 0.9604 | -0.0404 | 0.0638 | (0.8474, 1.0884) | 0.5267 |
| Diabetes mellitus (N=1366) | LV | 0.1084 | 0.1035 | 0.0366 | (0.0317, 0.1752) | **4.71x10<sup>-3\*\*</sup>** |
| Non-insulin-dependent diabetes mellitus (N=1055) |  | 0.1028 | 0.0970 | 0.0412 | (0.0162, 0.1778) | 0.0186<sup>†</sup> |
| Unspecified diabetes mellitus (N=913) |  | 0.1387 | 0.1308 | 0.0442 | (0.0440, 0.2175) | **0.0032\*\*** |
| Diabetes mellitus (N=1366) | 3rdV | 0.3030 | 0.3159 | 0.0400 | (0.2375, 0.3942) | **3.89x10<sup>-15\*\*</sup>** |
| Insulin-dependent diabetes mellitus (N=130) |  | 0.3270 | 0.3272 | 0.1268 | (0.0775, 0.5770) | **0.0104\*\*** |
| Non-insulin-dependent diabetes mellitus (N=1055) |  | 0.2875 | 0.2959 | 0.0449 | (0.2078, 0.3840) | **5.68x10<sup>-11\*\*</sup>** |
| Unspecified diabetes mellitus (N=913) |  | 0.3391 | 0.3523 | 0.0488 | (0.2566, 0.4480) | **7.47x10<sup>-13\*\*</sup>** |
| Diabetes mellitus (N=1366) | 4thV | 1.1326 | 0.1245 | 0.0829 | (0.9629, 1.3324) | 0.1328 |
| Disorders of other endocrine glands (N=336) | LV | 0.0247 | 0.0256 | 0.0807 | (-0.1329, 0.1842) | 0.7508 |
|  | 3rdV | 0.1749 | 0.1832 | 0.0815 | (0.0232, 0.3433) | 0.0249<sup>†</sup> |
|  | 4thV | 1.0126 | 0.0125 | 0.1790 | (0.7127, 1.4385) | 0.9443 |
| Other nutritional deficiencies (N=222) | LV | 0.0012 | 0.0013 | 0.1032 | (-0.2015, 0.2041) | 0.9898 |
|  | 3rdV | -0.0179 | -0.0187 | 0.1003 | (-0.2158, 0.1783) | 0.8520 |
|  | 4thV | 1.0916 | 0.0876 | 0.2128 | (0.7194, 1.6586) | 0.6806 |
| Obesity and other hyperalimentation (N=870) | LV | 0.1219 | 0.1185 | 0.0468 | (0.0268, 0.2102) | **0.0114\*\*** |
| Obesity (N=867) |  | 0.1151 | 0.1117 | 0.0468 | (0.0200, 0.2034) | 0.0170<sup>†</sup> |
| Obesity and other hyperalimentation (N=870) | 3rdV | 0.1743 | 0.1846 | 0.0510 | (0.0847, 0.2845) | **2.99x10<sup>-4\*\*</sup>** |
| Obesity (N=867) |  | 0.1706 | 0.1808 | 0.0511 | (0.0807, 0.2810) | **4.08x10<sup>-4\*\*</sup>** |
| Obesity and other hyperalimentation (N=870) | 4thV | 1.1500 | 0.1397 | 0.1060 | (0.9343, 1.4159) | 0.1875 |
| Metabolic disorders (N=5840) | LV | 0.0384 | 0.0376 | 0.0181 | (0.0021, 0.0732) | 0.0381<sup>†</sup> |
| Disorders of lipoprotein metabolism and other lipidaemias (N=5435) |  | 0.0529 | 0.0516 | 0.0187 | (0.0149, 0.0883) | **0.0059\*\*** |
| Metabolic disorders (N=5840) | 3rdV | 0.1256 | 0.1273 | 0.0188 | (0.0905, 0.1641) | **1.21x10<sup>-11\*\*</sup>** |
| Disorders of lipoprotein metabolism and other lipidaemias (N=5435) |  | 0.1379 | 0.1391 | 0.0194 | (0.1011, 0.1770) | **7.13x10<sup>-13\*\*</sup>** |
| Volume depletion (N=168) |  | 0.3331 | 0.3528 | 0.1175 | (0.1217, 0.5840) | **0.0029\*\*** |
| Metabolic disorders (N=5840) | 4thV | 1.0569 | 0.0554 | 0.0400 | (0.9772, 1.1431) | 0.1664 |

\*\*Uncorrected $p \leq$ critical FDR threshold \*Uncorrected $p \leq 0.05$

#### 3.2.2 Mental and behavioral disorders

Within ICD-10 chapter V, the strongest associations were with substance use (Table 5). Mental and behavioural disorders due to psychoactive substance use were associated with greater CPcal in the LV (d = 0.1521, p = 0.0032) and the 3rdV (d = 0.2680, p = 2.20×10^−7^), an effect that was most significant for tobacco use disorders, with greater CPcal in the LV (d = 0.2049, p = 8.58×10^−4^) and the 3rdV (d = 0.3142, p = 3.50×10^−7^). Mood (affective) disorders were associated with greater CPcal in the LV (d = 0.0881, p = 4.94×10^−4^) and the 3rdV (d = 0.0913, p = 3.03×10^−4^), with a similar effect for depressive episodes in the LV (d = 0.0856, p = 7.47×10^−4^) and the 3rdV (d = 0.0950, p = 1.84×10^−4^). Neurotic, stress-related and somatoform disorders were associated with greater CPcal in the LV (d = 0.1365, p = 9.06×10^−4^) and the 3rdV (d = 0.1712, p = 3.19×10^−5^). However, the LV association is significant after FDR correction. Within this level, other anxiety disorders were associated with greater CPcal in the LV (d = 0.1538, p = 0.0048) and the 3rdV (d = 0.1881, p = 5.64×10^−4^). Mental disorders and behavioral syndromes associated with physiological disturbances were not significantly associated with CPcal in any ventricle.

**Table 5.** Associations between choroid plexus calcification (CPcal) volumes and level-1 diagnostic categories within ICD-10 chapter V (Mental and behavioral disorders). Significant level-1 categories are further broken down into specific ICD-10 disorders. Linear regression models were used for CPcal in the lateral ventricles (LV) and third ventricle (3rdV), and logistic regression for the fourth ventricle (4thV) due to its binary distribution (calcified vs not calcified).

| ICD-10 chapters | CPcal Region | Effect Size | β | SE | 95% CI | P-value |
| --- | --- | --- | --- | --- | --- | --- |
| Organic, including symptomatic, mental disorders (N=119) | LV | 0.2417 | 0.2213 | 0.1215 | (-0.0182, 0.4607) | 0.0699 |
|  | 3rdV | 0.1694 | 0.1612 | 0.1263 | (-0.0878, 0.4101) | 0.2033 |
|  | 4thV | 0.7003 | -0.3562 | 0.2960 | (0.3893, 1.2457) | 0.2287 |
| Mental and behavioural disorders due to psychoactive substance use (N=760) | LV | 0.1521 | 0.1529 | 0.0518 | (0.0514, 0.2545) | **0.0032\*\*** |
| Mental and behavioural disorders due to use of tobacco (N=538) |  | 0.2049 | 0.2066 | 0.0618 | (0.0853, 0.3278) | **8.58x10<sup>-4</sup>\*\*** |
| Mental and behavioural disorders due to psychoactive substance use (N=760) | 3rdV | 0.2680 | 0.2654 | 0.0510 | (0.1654, 0.3654) | **2.20x10<sup>-7</sup>\*\*** |
| Mental and behavioural disorders due to use of tobacco (N=538) |  | 0.3142 | 0.3171 | 0.0619 | (0.1958, 0.4385) | **3.50x10<sup>-7</sup>\*\*** |
| Mental and behavioural disorders due to psychoactive substance use (N=760) | 4thV | 1.1574 | 0.1462 | 0.1111 | (0.9309, 1.4395) | 0.1885 |
| Mood [affective] disorders (N=3138) | LV | 0.0881 | 0.0923 | 0.0265 | (0.0404, 0.1441) | **4.94x10<sup>-4</sup>\*\*** |
| Depressive episode (N=3112) |  | 0.0856 | 0.0898 | 0.0266 | (0.0376, 0.1419) | **7.47x10<sup>-4</sup>\*\*** |
| Mood [affective] disorders (N=3138) | 3rdV | 0.0913 | 0.0997 | 0.0276 | (0.0456, 0.1537) | **3.03x10<sup>-4</sup>\*\*** |
| Depressive episode (N=3112) |  | 0.0950 | 0.1037 | 0.0277 | (0.0494, 0.1580) | **1.84x10<sup>-4</sup>\*\*** |
| Mood [affective] disorders (N=3138) | 4thV | 1.0494 | 0.0483 | 0.0560 | (0.9404, 1.1712) | 0.3887 |
| Neurotic, stress-related and somatoform disorders (N=1190) | LV | 0.1365 | 0.1417 | 0.0427 | (0.0581, 0.2253) | **9.06x10<sup>-4</sup>\*\*** |
| Other anxiety disorders (N=681) |  | 0.1538 | 0.1614 | 0.0571 | (0.0494, 0.2735) | **0.0048\*\*** |
| Neurotic, stress-related and somatoform disorders (N=1190) | 3rdV | 0.1712 | 0.1777 | 0.0426 | (0.0941, 0.2613) | **3.19x10<sup>-5</sup>\*\*** |
| Other anxiety disorders (N=681) |  | 0.1881 | 0.1979 | 0.0573 | (0.0856, 0.3103) | **5.64x10<sup>-4</sup>\*\*** |
| Neurotic, stress-related and somatoform disorders (N=1190) | 4thV | 1.0956 | 0.0913 | 0.0904 | (0.9176, 1.3082) | 0.3129 |
| Behavioural syndromes associated with physiological disturbances and physical factors (N=226) | LV | 0.0943 | 0.1090 | 0.1100 | (-0.1072, 0.3251) | 0.3223 |
|  | 3rdV | 0.1920 | 0.2192 | 0.1086 | (0.0057, 0.4327) | 0.0442* |
|  | 4thV | 1.1386 | 0.1298 | 0.2106 | (0.7535, 1.7219) | 0.5375 |

\*\*Uncorrected p ≤ critical FDR threshold \*Uncorrected p ≤ 0.05

#### 3.2.3 Diseases of the nervous system

Within ICD-10 chapter VI, significant associations were largely confined to the 3rdV (Table 6). Episodic and paroxysmal disorders were associated with greater CPcal in the 3rdV (d = 0.0904, p = 4.16×10^−4^), as were sleep disorders (d = 0.1928, p = 0.0045). Migraine showed an association with 3rdV that did not survive FDR correction (d = 0.0829, p = 0.0070). Nerve, nerve root and plexus disorders were associated with greater CPcal in the LV (d = 0.1041, p = 0.0053) and the 3rdV (d = 0.0996, p = 0.0077). At the disorder level, mononeuropathies of the upper limb were associated with greater CPcal in the LV (d = 0.1485, p = 0.0032) only; the 3rdV association was not significant. Demyelinating diseases of the central nervous system showed an association with 3rdV that did not survive FDR correction (d = 0.2777, p = 0.0290). No significant associations were found for inflammatory diseases of the central nervous system, extrapyramidal and movement disorders, or polyneuropathies.

**Table 6.**
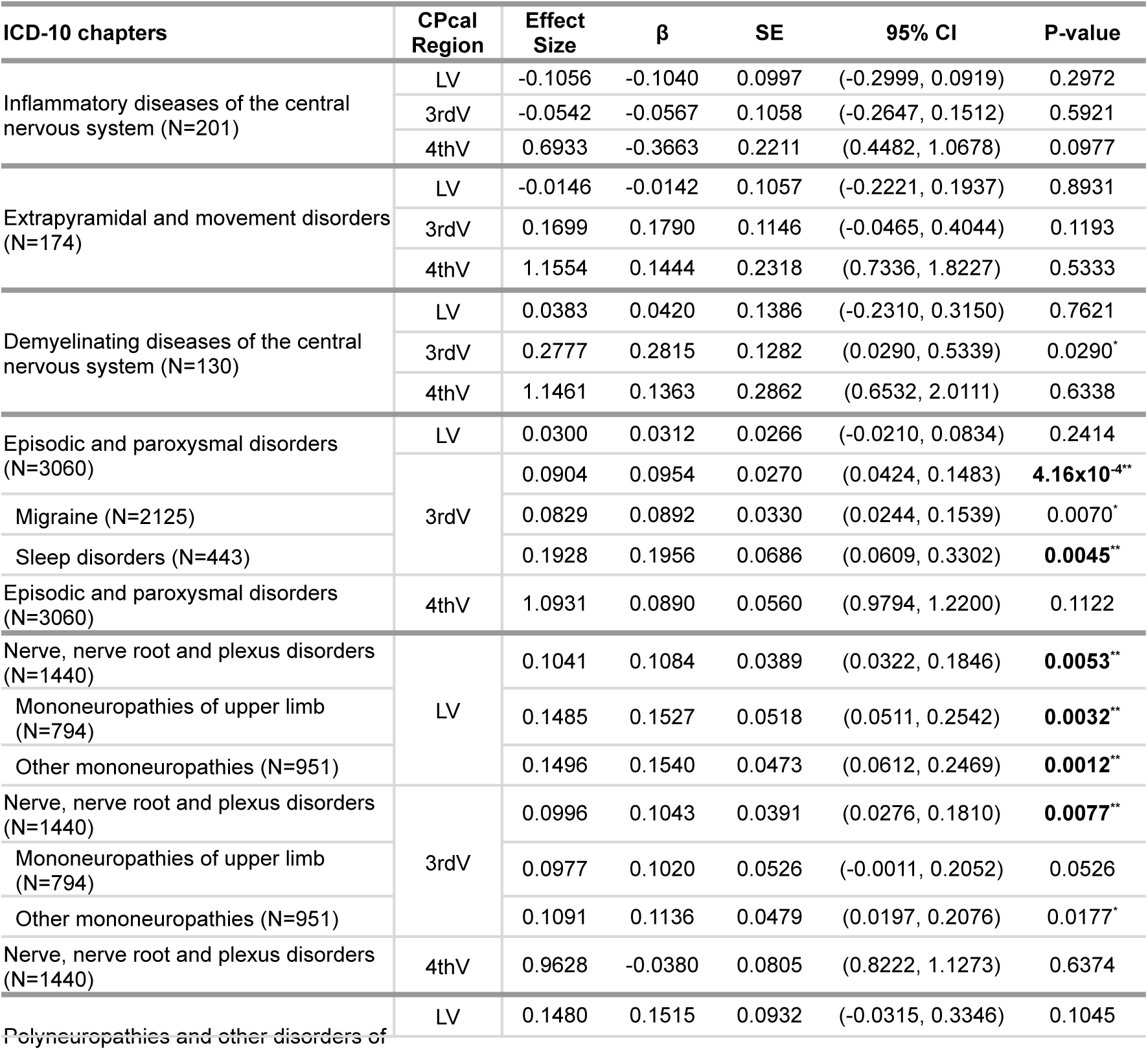
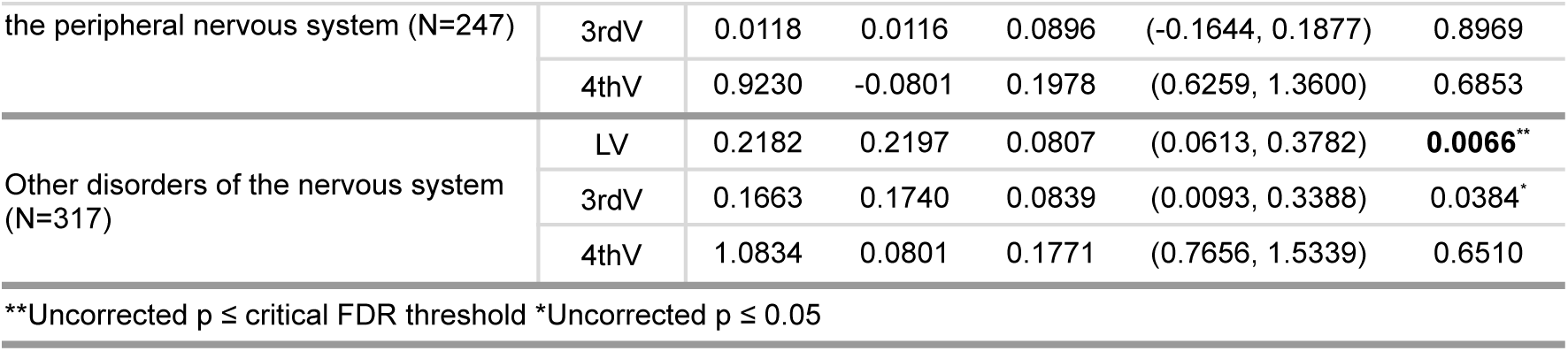
Associations between choroid plexus calcification (CPcal) volumes and level-1 diagnostic categories within ICD-10 chapter VI (Diseases of the nervous system). Significant level-1 categories are further broken down into specific ICD-10 disorders. Linear regression models were used for CPcal in the lateral ventricles (LV) and third ventricle (3rdV), and logistic regression for the fourth ventricle (4thV) due to its binary distribution (calcified vs not calcified).

#### 3.2.4 Diseases of the circulatory system

Within ICD-10 chapter IX, cardiovascular conditions were associated with larger CPcal, predominantly in the 3rdV (Table 7). Hypertensive diseases were associated with greater CPcal in the LV (d = 0.0547, p = 0.0060) and the 3rdV (d = 0.1402, p = 2.00×10^−12^), and essential (primary) hypertension in the 3rdV (d = 0.1430, p = 7.26×10^−13^). Ischaemic heart disease was associated with greater CPcal in the 3rdV (d = 0.0809, p = 0.0125), although associations with acute myocardial infarction and chronic ischaemic heart disease did not survive FDR correction. Pulmonary heart disease and diseases of pulmonary circulation were associated with greater CPcal in the 3rdV (d = 0.2480, p = 0.0010). Pulmonary embolism showed a similar 3rdV effect (d = 0.2484, p = 0.0015). Other forms of heart disease were associated with greater CPcal in the 3rdV (d = 0.0861, p = 0.0030), particularly among individuals with atrial fibrillation and flutter (d = 0.1263, p = 0.0029) and heart failure (d = 0.1785, p = 0.0350). Cerebrovascular disease was associated with greater CPcal in the 3rdV (d = 0.1656, p = 0.0033).

**Table 7.**
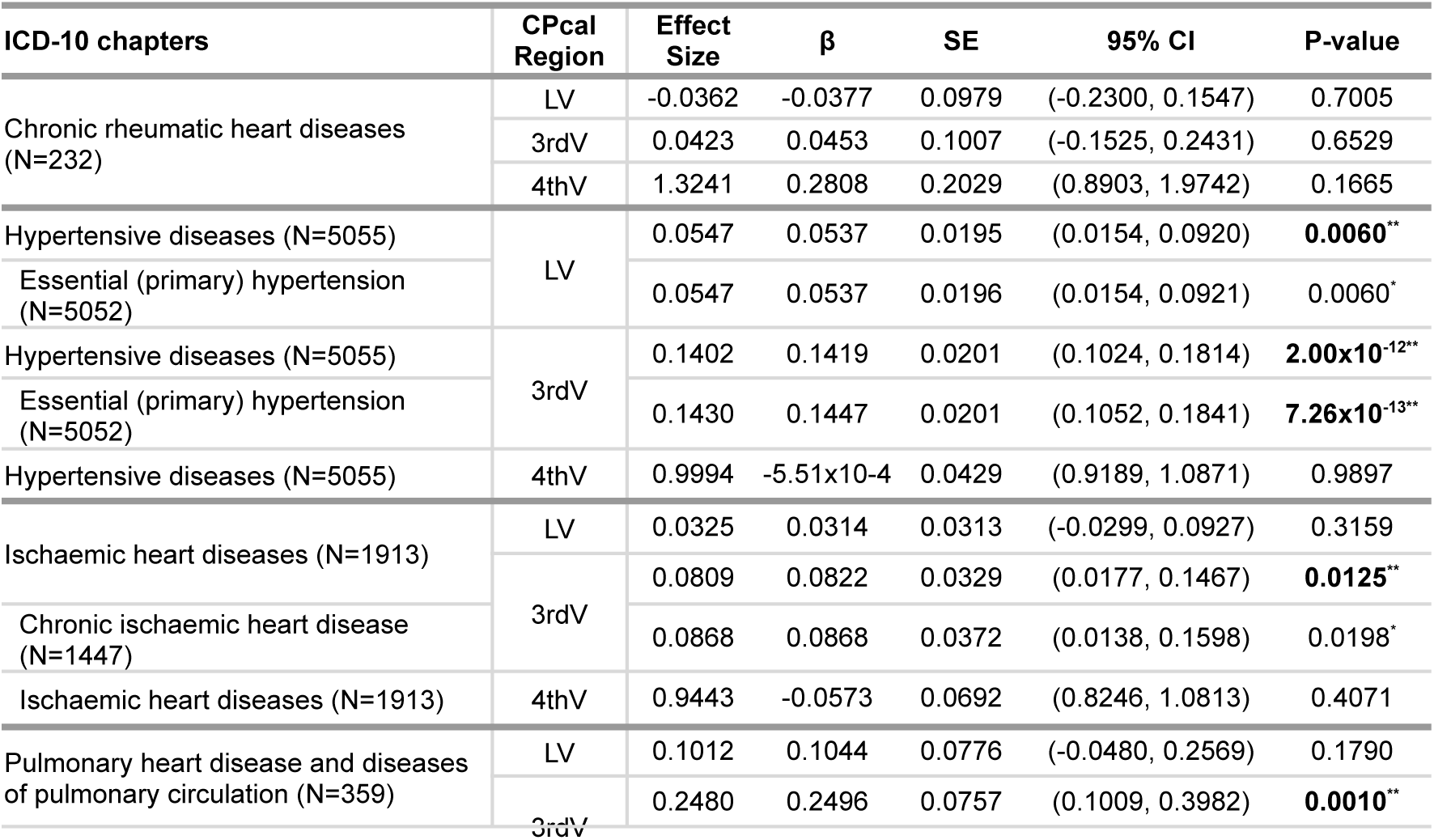
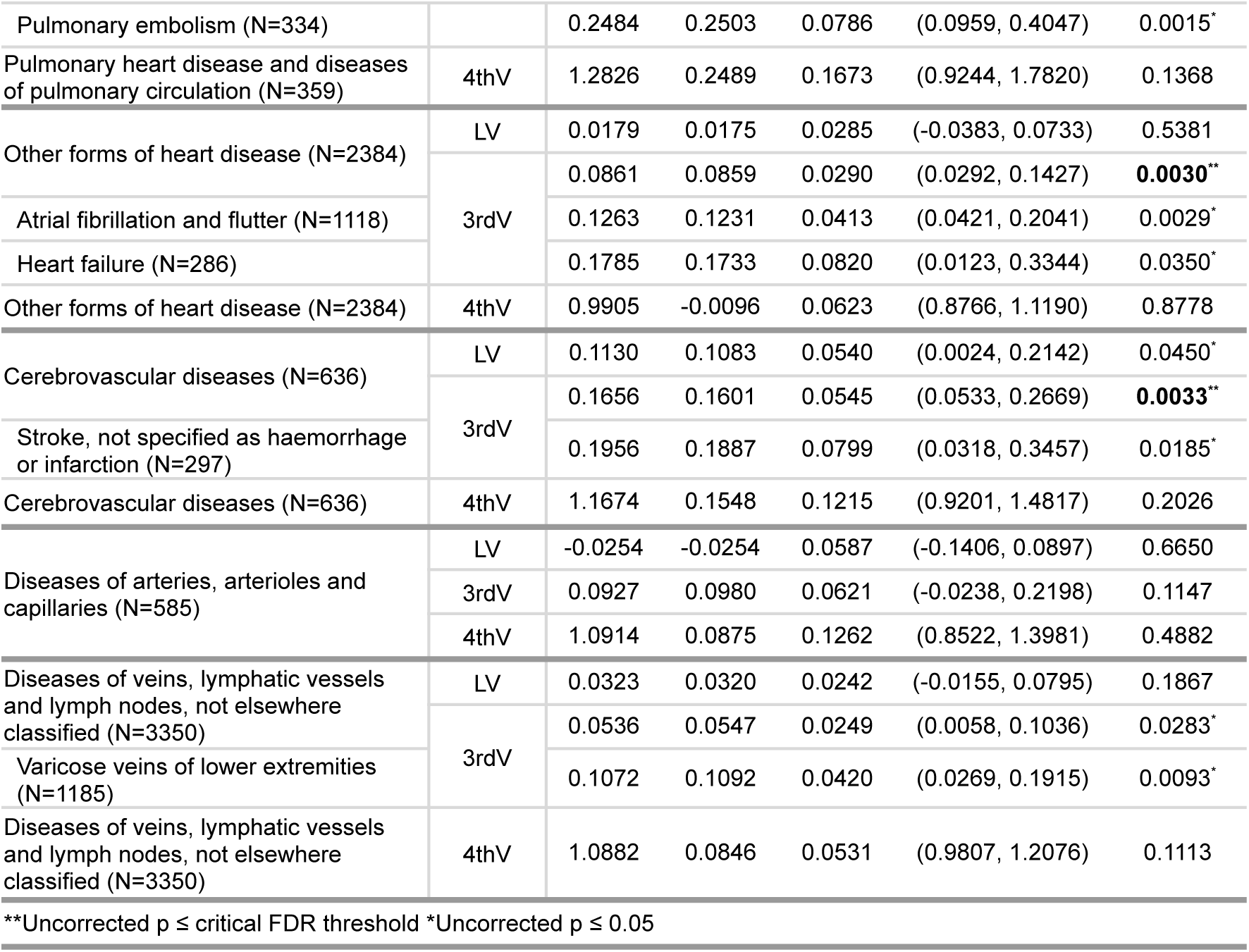
Associations between choroid plexus calcification (CPcal) volumes and level-1 diagnostic categories within ICD-10 chapter IX (Diseases of the circulatory system). Significant level-1 categories are further broken down into specific ICD-10 disorders. Linear regression models were used for CPcal in the lateral ventricles (LV) and third ventricle (3rdV), and logistic regression for the fourth ventricle (4thV) due to its binary distribution (calcified vs not calcified).

In the exploratory parental-history proxy analysis, maternal ADRD history was associated with increased lateral-ventricle CPcal (d = 0.0631, p = 0.0033; Table 8). Paternal history (d = −0.0306, p = 0.30) and either parent history (d = 0.0278, p = 0.12) were not associated with lateral ventricle CPcal, and no significant associations were observed in the third or fourth ventricles.

**Table 8.** Associations between choroid plexus calcification (CPcal) volumes and family history of Alzheimer’s disease or related dementias (ADRD). Linear regression models were used for CPcal in the lateral ventricles (LV) and third ventricle (3rdV), and logistic regression for the fourth ventricle (4thV) due to its binary distribution (calcified vs not calcified).

| Family history of ADRD | CPcal Region | Effect Size | β | SE | 95% CI | P-value |
| --- | --- | --- | --- | --- | --- | --- |
| Either parents | LV | 0.0278 | 0.0279 | 0.0180 | (-0.0075, 0.0632) | 0.1221 |
|  | 3rdV | 0.0018 | 0.0018 | 0.0184 | (-0.0341, 0.0378) | 0.9204 |
|  | 4thV | 1.0233 | 0.0230 | 0.0389 | (0.9481, 1.1044) | 0.5540 |
| Both parents | LV | -0.0328 | -0.0353 | 0.0717 | (-0.1761, 0.1054) | 0.6225 |
|  | 3rdV | -0.1418 | -0.1557 | 0.0731 | (-0.2992, -0.0122) | 0.0335<sup>*</sup> |
|  | 4thV | 1.0955 | 0.0912 | 0.1517 | (0.8139, 1.4759) | 0.5476 |
| Mother | LV | 0.0631 | 0.0633 | 0.0215 | (0.0211, 0.1055) | **0.0033<sup>\*\*</sup>** |
|  | 3rdV | 0.0225 | 0.0232 | 0.0222 | (-0.0202, 0.0667) | 0.2942 |
|  | 4thV | 1.0346 | 0.0340 | 0.0466 | (0.9444, 1.1335) | 0.4647 |
| Father | LV | -0.0306 | -0.0317 | 0.0305 | (-0.0915, 0.0282) | 0.2998 |
|  | 3rdV | 0.0019 | 0.0020 | 0.0308 | (-0.0584, 0.0625) | 0.9478 |
|  | 4thV | 1.0171 | 0.0169 | 0.0648 | (0.8957, 1.1549) | 0.7940 |

\*\*Uncorrected $p \leq$ critical FDR threshold \*Uncorrected $p \leq 0.05$

### 3.3 Disease progression

In a subset of participants first diagnosed after their baseline imaging visit, CPcal volumes derived from the scan before diagnosis were compared with age and sex-matched controls who did not develop the disease during follow-up imaging (Table 9). Participants who developed endocrine and metabolic disease had greater CPcal in the LV (d = 0.1183, p = 9.30×10^−5^) and the 3rdV (d = 0.1413, p = 3.08×10^−6^) than those who did not develop the disease. Similarly, participants who developed mental and behavioral disorders had greater CPcal in the LV (d = 0.1794, p = 3.40×10^−4^) and the 3rdV (d = 0.1649, p = 9.86×10^−4^). Also, participants who developed circulatory-system disease had greater CPcal in the LV (d = 0.0805, p = 0.0023) and the 3rdV (d = 0.1273, p = 1.52×10^−6^). Participants who developed nervous-system disease had greater CPcal in the 3rdV (d = 0.1105, p = 0.0253) at baseline compared to controls, but no significant effects were found in the LV. Associations with individual level-1 diagnoses within each chapter are reported in the Supplementary Material (Tables S2–S5).

**Table 9.**
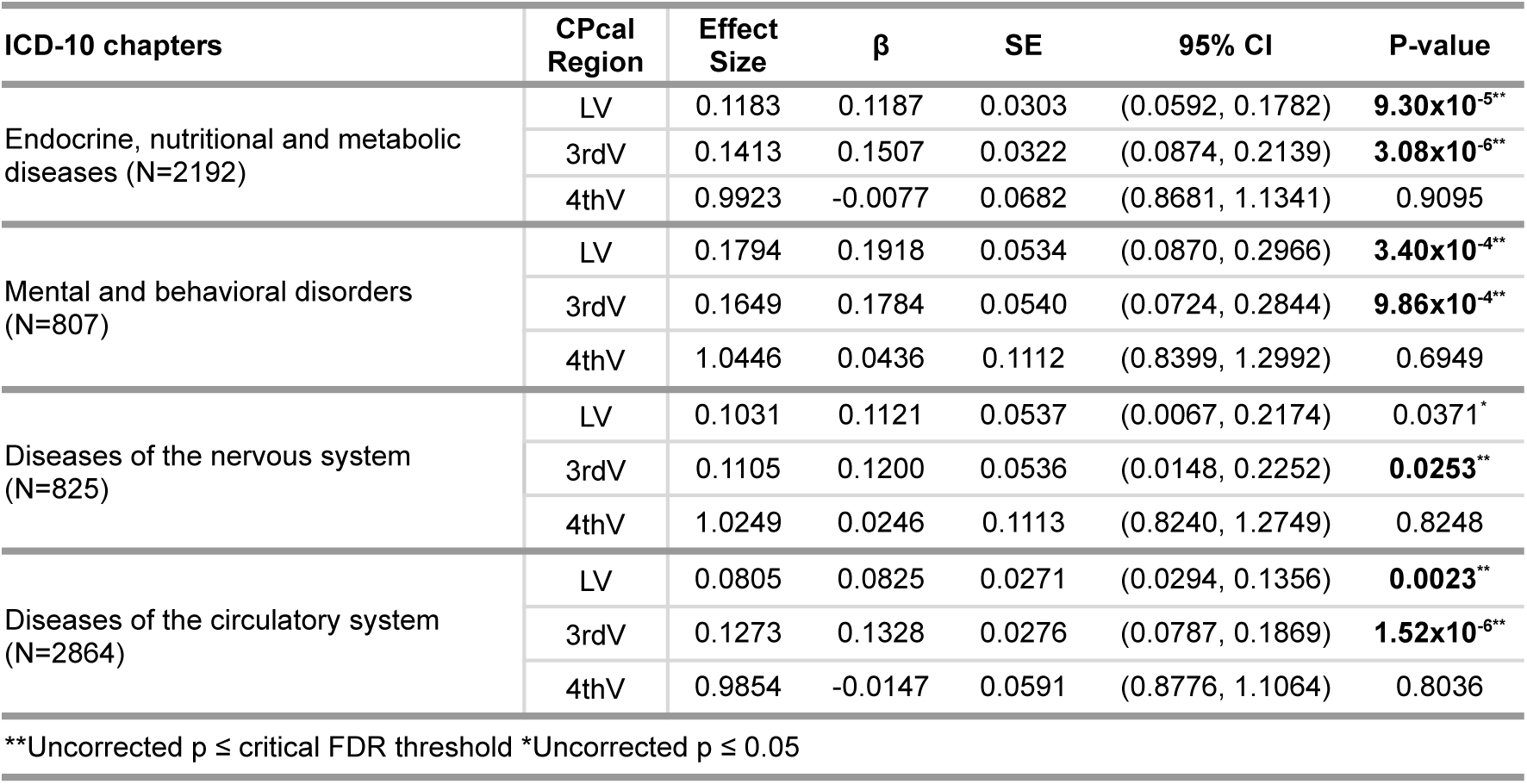
Associations between choroid plexus calcification (CPcal) volumes and ICD-10 diagnostic chapters, focusing on endocrine and metabolic disorders, mental and behavioral disorders, diseases of the nervous system, and diseases of the circulatory system. Linear regression models were used for CPcal in the lateral ventricles (LV) and third ventricle (3rdV), and logistic regression for the fourth ventricle (4thV) due to its binary distribution (calcified vs not calcified).

### 3.4 Cardiovascular risk factor associations with CPcal volumes

Given the associations with circulatory disease, we tested cardiovascular and metabolic risk factors measured at the imaging visit (Table 10). Larger CPcal volumes were associated with higher body mass index, whole-body fat mass, basal metabolic rate, waist-to-hip ratio, pulse rate, and cardiac output. The largest effects were in the 3rdV (e.g., body mass index r = 0.1729, p = 8.87×10^−185^).

**Table 10.** Associations between choroid plexus calcification (CPcal) volumes and cardiovascular risk factors. Linear regression models were used for CPcal in the lateral ventricles (LV) and third ventricle (3rdV), and logistic regression for the fourth ventricle (4thV) due to its binary distribution (calcified vs not calcified).

| Cardiovascular Risk Factors | CPcal Region | Effect Size | $\beta$ | SE | 95% CI | P-value |
| --- | --- | --- | --- | --- | --- | --- |
| Pulse Rate | LV | 0.0196 | 0.0018 | $5.78 \times 10^{-4}$ | ($6.37 \times 10^{-4}$, 0.0029) | **0.0022\*\*** |
| | 3rdV | 0.0788 | 0.0072 | $5.86 \times 10^{-4}$ | (0.0061, 0.0084) | **7.46x10<sup>-35\*\*</sup>** |
|  | 4thV | 1.0021 | 0.0021 | 0.0012 | (0.9997, 1.0045) | 0.0873 |
| Cardiac Output | LV | 0.0229 | 0.0159 | 0.0044 | (0.0072, 0.0246) | **3.44x10<sup>-4\*\*</sup>** |
|  | 3rdV | 0.0481 | 0.0341 | 0.0045 | (0.0252, 0.0430) | **5.72x10<sup>-14\*\*</sup>** |
|  | 4thV | 1.0041 | 0.0041 | 0.0093 | (0.9855, 1.0225) | 0.6575 |
| Basal Metabolic Rate (BMR) | LV | 0.0853 | $1.17 \times 10^{-4}$ | $8.20 \times 10^{-6}$ | ($1.01 \times 10^{-4}$, $1.33 \times 10^{-4}$) | **6.18x10<sup>-46\*\*</sup>** |
| | 3rdV | 0.1332 | $1.86 \times 10^{-4}$ | $8.33 \times 10^{-6}$ | ($1.70 \times 10^{-4}$, $2.03 \times 10^{-4}$) | **7.60x10<sup>-110\*\*</sup>** |
| | 4thV | 1.0000 | $4.40 \times 10^{-5}$ | $1.74 \times 10^{-5}$ | (1.0000, 1.0001) | **0.0114\*\*** |
| Whole Body Fat Mass (BFM) | LV | 0.0535 | 0.0068 | $7.61 \times 10^{-4}$ | (0.0053, 0.0083) | **5.03x10<sup>-19\*\*</sup>** |
| | 3rdV | 0.1804 | 0.0233 | $7.65 \times 10^{-4}$ | (0.0218, 0.0248) | **4.03x10<sup>-201\*\*</sup>** |
|  | 4thV | 1.0044 | 0.0044 | 0.0016 | (1.0011, 1.0076) | **0.0078\*\*** |
| Waist-to-Hip Ratio (WHR) | LV | 0.0352 | 0.5583 | 0.0943 | (0.3734, 0.7432) | **3.28x10<sup>-9\*\*</sup>** |
|  | 3rdV | 0.1361 | 2.2050 | 0.0954 | (2.0179, 2.3920) | **5.12x10<sup>-117\*\*</sup>** |
|  | 4thV | 2.3573 | 0.8575 | 0.2021 | (1.5862, 3.5029) | **2.21x10<sup>-5\*\*</sup>** |
| Body Mass Index (BMI) | LV | 0.0627 | 0.0157 | 0.0015 | (0.0128, 0.0187) | **1.50x10<sup>-25\*\*</sup>** |
|  | 3rdV | 0.1729 | 0.0442 | 0.0015 | (0.0413, 0.0472) | **8.87x10<sup>-185\*\*</sup>** |
|  | 4thV | 1.0108 | 0.0107 | 0.0032 | (1.0044, 1.0172) | **9.19x10<sup>-4\*\*</sup>** |

\*\*Uncorrected $p \leq$ critical FDR threshold \*Uncorrected $p \leq 0.05$

### 3.5 Brain Volume and Microstructural Associations with CPcal Volume

#### 3.5.1 Cross-sectional: Cortical, Subcortical regions

Across all participants, larger CPcal volume in the lateral ventricles (LV) was associated with lower cortical thickness and greater cortical surface area globally (Figure 2). Among subcortical volumes, larger LV CPcal was associated with greater caudate (r = 0.0462, p = 3.65×10^−15^) volume, putamen (r = 0.0325, p = 3.05×10^−8^) volume, hippocampus (r = 0.0352, p = 2.07×10^−9^) volume, and accumbens (r = 0.0362, p = 6.61×10^−10^) volume.Larger CPcal volume in the third ventricle (3rdV) was associated with predominantly greater cortical thickness and greater cortical surface area (Figure 2). Among subcortical volumes, larger 3rdV CPcal was associated with greater caudate (r = 0.1077, p = 1.36×10^−75^) volume, putamen (r = 0.0611, p = 2.17×10^−25^) volume, and accumbens (r = 0.0630, p = 5.93×10^−27^) volume, and with lower pallidum (r = −0.0590, p = 8.32×10^−24^) volume and hippocampus (r = −0.0406, p = 4.58×10^−12^) volume. The presence of CPcal in the fourth ventricle (4thV) was associated with greater cortical thickness and lower cortical surface area, in a smaller number of regions (Figure 2). Among subcortical volumes, 4thV CPcal was associated with greater caudate (d = 0.0510, p = 4.13×10^−5^) volume and putamen (d = 0.0430, p = 5.52×10^−4^) volume, and with lower accumbens (d = −0.0315, p = 0.0114) volume.

**Figure 2.**
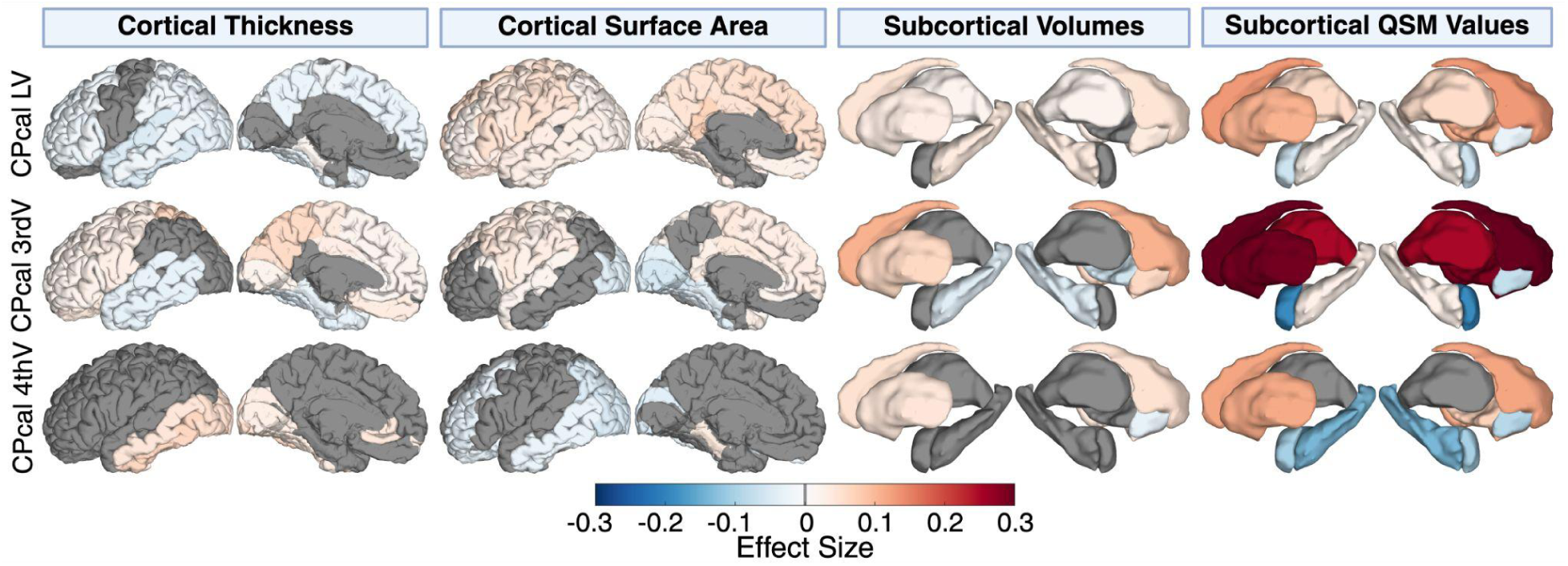
Associations between choroid plexus calcification (CPcal) volumes, and cortical (thickness and surface area) subcortical volumes regions and quantitative susceptibility mapping (QSM) values in subcortical regions. Linear regression models were used for CPcal in the lateral ventricles (LV), third ventricle (3rdV) and fourth ventricle (4thV).

#### 3.5.2 Cross-sectional Microstructure: subcortical QSM values and WM dMRI

##### 3.5.2.1 Quantitative Susceptibility Mapping (QSM) values

Larger CPcal volume in the LV was associated with higher magnetic susceptibility (QSM) in the thalamus (r = 0.0571, p = 2.04×10^−22^), caudate (r = 0.1310, p = 2.92×10^−111^), putamen (r = 0.1066, p = 3.99×10^−74^), pallidum (r = 0.1099, p = 1.07×10^−78^), and hippocampus (r = 0.0228, p = 1.05×10^−4^), and with lower susceptibility in the amygdala (r = −0.0635, p = 2.60×10^−27^) and accumbens (r = −0.0303, p = 2.40×10^−7^).

##### 3.5.2.2 Diffusion measures in white matter tracts

Larger CPcal volume in the LV was associated with white matter microstructure changes (Figure 3). Fractional anisotropy (FA) was higher in 11 tracts (4 projection, 3 brainstem, 2 association, 2 commissural) and lower in the posterior corona radiata (r = −0.0166, p = 0.0056), fornix (r = −0.0423, p = 1.39×10^−12^), and fornix (cres) / stria terminalis (r = −0.0239, p = 6.12×10^−5^). Mean diffusivity (MD) was lower in 13 tracts (3 projection, 6 brainstem, 4 association) and higher in the fornix (r = 0.0269, p = 6.82×10^−6^).

**Figure 3.**
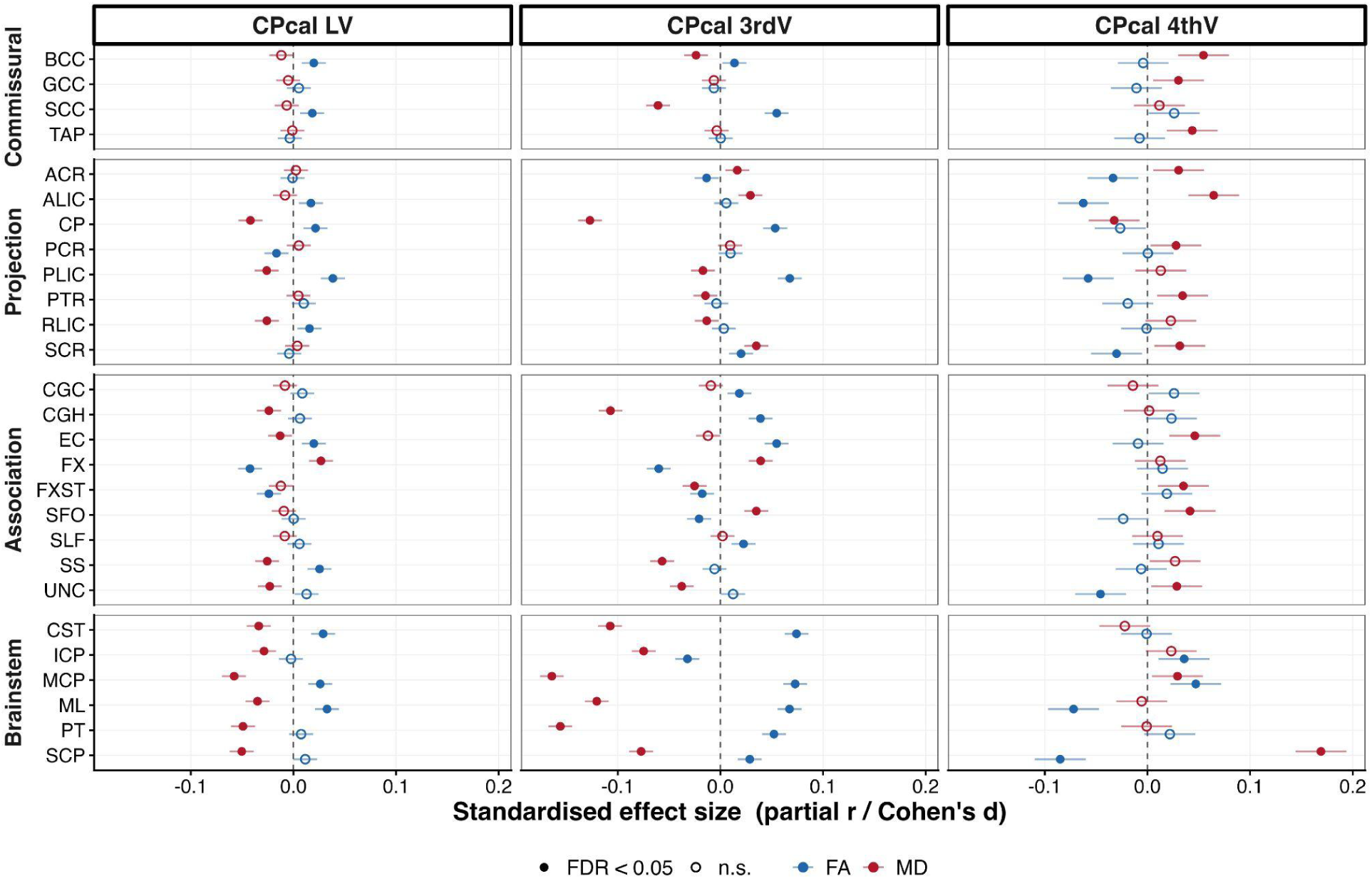
Associations between choroid plexus calcification (CPcal) volumes and diffusion measures (fractional anisotropy and mean diffusivity) of white matter tracts, defined based on JHU-ICBM template. Linear regression models were used for CPcal in the lateral ventricles (LV), third ventricle (3rdV) and fourth ventricle (4thV). White matter bundles are grouped by tract category (commissural, projection, association, and brain stem). Fractional anisotropy (FA) is shown in blue and mean diffusivity (MD) in red. The filled circle indicates associations surviving false discovery rate (FDR) correction (P_FDR_ < 0.05). The abbreviations of the white matter bundles are as follows: BCC, body of corpus callosum; GCC, genu of corpus callosum; SCC, splenium of corpus callosum; TAP, tapetum; ACR, anterior corona radiata; ALIC, anterior limb of internal capsule; CP, cerebral peduncle; PCR, posterior corona radiata; PLIC, posterior limb of internal capsule; PTR, posterior thalamic radiation; RLIC, retrolenticular part of internal capsule; SCR, superior corona radiata; CGC, cingulum (cingulate gyrus); CGH, cingulum (hippocampal portion); EC, external capsule; FX, fornix (column and body); FXST, fornix cres/stria terminalis; SFO, superior fronto-occipital fasciculus; SLF, superior longitudinal fasciculus; SS, sagittal stratum; UNC, uncinate fasciculus; CST, corticospinal tract; ICP, inferior cerebellar peduncle; MCP, middle cerebellar peduncle; ML, medial lemniscus; PT, pontine crossing tract; SCP, superior cerebellar peduncle.

## 4 Discussion

Our study aimed to characterize the extent to which CPcal in the lateral, third, and fourth ventricles as determined by MRI-derived quantitative susceptibility mapping (QSM) can serve as a biomarker for a variety of conditions. Specifically, we assessed associations with age, sex, alterations in brain region volumes, endocrine, metabolic, circulatory and neuropsychiatric diseases. We found: (1) a positive association between the CPcal and age; (2) larger CPcal volumes were associated with endocrine, nutritional, and metabolic diseases; mental and behavioral disorders; and diseases of the nervous and circulatory systems; and (3) larger CPcal volume in the lateral ventricles was associated with lower cortical thickness, greater cortical surface area, and larger subcortical volumes. Most notably, the strongest associations were identified with the calcification extent in the CP of the third ventricle, an often ignored region for assessing the choroid plexus.

The CP undergoes a range of changes with aging, and calcification is one of the earliest and most consistent structural markers of that process [Butler et al., 2023]. In our study, CPcal and volume were only weakly correlated (r = 0.22; Section 3.1), suggesting that these measures capture distinct aspects of CP biology: calcification reflects mineral deposition, and volume reflects irregular fibrotic changes in the stroma and basement membrane thickening [Hidaka et al., 2024]. This distinction may be biologically meaningful. In a previous study, CPcal, but not CP volume, independently predicted cortical microglial activation on TSPO PET [Butler et al., 2023]. Moreover, T1-based segmentation pipelines of CP may incompletely capture calcification [Tadayon et al., 2020], suggesting that volumetric measures may not fully capture calcification-related changes. Two additional findings support this distinction. First, CPcal increased with age and was greater in males, consistent with large CT studies showing that CPcal is common after midlife, predominantly occurs in the atria of the lateral ventricles, and is more prevalent in males [Bhatt, 2022; Saade et al., 2019; Yalcin et al., 2016]. Second, a longitudinal study on CP volume showed a different trend: volume increased at more than twice the annual rate in females than in males [Novakova Martinkova et al., 2023]. The contrasting sex results, greater volumetric enlargement in females but greater calcification in males, further suggest that these measures are not interchangeable and may capture distinct aspects of CP pathology. In the same cohort, a prospective analysis of 45,306 UK Biobank participants found that CP volume increased with age and was associated with incident all-cause dementia; Mendelian randomization supported a causal contribution [Yu et al., 2026]. That study derived CP volume and mean T1 signal intensity segmented using FreeSurfer, interpreting T1 intensity as a marker of tissue water content and microstructural integrity. Although calcification was observed among dementia-related CP changes, neither measure directly quantified mineral deposition, and the CP was limited to the lateral ventricles [Yu et al., 2026]. QSM directly quantifies the mineral component and extends its assessment to the third and fourth ventricles.

A consistent feature of our findings was their regional specificity. Across metabolic, psychiatric, neurological, and circulatory disorders, associations with CPcal were strongest in the third ventricle for both prevalent and incident diagnoses. This finding is notable as CP in the third ventricle has received little attention in imaging studies, which have focused mainly on the lateral ventricles. Its distinct biological features may partly explain this regional difference. The third ventricle CP is located in the diencephalic roof, adjacent to hypothalamic and circumventricular structures with fenestrated capillaries and direct access to systemic signals [Kaiser and Bryja, 2020; Municio et al., 2023]. It is also a major source of apolipoprotein E in CSF [Municio et al., 2023], while third ventricle tanycytes have been implicated in iron exchange between CSF and brain tissue [Ficiarà et al., 2022]. Age-related loss of epithelial microvilli has been reported in the CP of the lateral and third ventricles, but not in the fourth [Gonzalez-Marrero et al., 2022], consistent with our finding that fourth-ventricle associations were weaker and largely null. Given its location at the interface between systemic circulation and hypothalamic metabolic regulation, the third-ventricle CP may be particularly sensitive to systemic metabolic and vascular burdens, as suggested by our findings.

Across the four ICD-10 disease categories examined - metabolic, psychiatric, neurological, and circulatory disorders - our findings converge on two broader themes. The first is systemic inflammatory and metabolic burden. Greater CPcal was associated with diabetes, obesity, lipoprotein disorders, higher body mass index and fat mass, increased cardiac output, hypertensive, ischaemic, and cerebrovascular disease, and substance use. The CP represents a key interface between peripheral circulation and the CSF, and CP volume has similarly been associated with adiposity, diabetes, and white matter hyperintensity burden [Hidaka et al., 2024; Li et al., 2024; Ricigliano and Stankoff, 2023]. The second theme is dysregulated mineralization and ectopic calcification. Since CPcal reflects mineral deposition rather than tissue bulk, inflammatory enlargement alone is unlikely to fully explain these associations. Dysregulation of ectopic calcification may provide a more direct mechanism, consistent with reduced circulating levels of the calcification inhibitors osteopontin and fetuin-A in individuals with CPcal [Yevgi et al., 2022]. A vascular contribution is also possible: experimental forebrain ischaemia increases calcium influx across the CP and induces persistent endothelial injury in the CP of the lateral and third ventricles [Ikeda et al., 1992]. Our psychiatric findings further extend previous CT studies linking CPcal to depressive symptoms and psychotic illness severity[Demeestere et al., 2015; Lizano et al., 2019; Marinescu et al., 2013; Sandyk et al., 1990]. As these studies focused on the lateral ventricles, our findings highlight the potential importance of CP in the third ventricle. Particularly, associations were also present in participants diagnosed after imaging, indicating that elevated CPcal may be detectable before clinical diagnosis.

In our brain microstructural analyses, the associations similarly converged on a common pattern. Greater CPcal in the lateral ventricle was associated with lower cortical thickness and greater surface area, larger subcortical volumes, higher QSM value in iron rich deep grey matter nuclei, and higher FA with lower MD across most white matter tracts. Importantly, these associations were regionally specific: cortical thickness associations were mostly positive for CPcal in the third ventricle, contrasting with the lateral ventricle findings and indicating that reduced cortical thickness is not a general feature of CPcal. Across imaging modalities, the overall profile was more consistent with variation in iron-related susceptibility than with widespread tissue loss. CP psammoma bodies are dystrophic calcifications in which iron is commonly co-deposited [Alcolado et al., 1986]. The CP is also a major site of iron storage and transferrin secretion into the CSF [Ficiarà et al., 2022; Ward et al., 2014]. Consistent with these findings, the higher susceptibility observed in iron-rich nuclei aligns with the established sensitivity of QSM to non-heme iron [Ravanfar et al., 2021]. The white matter findings, characterized by higher FA and lower MD, do not typically indicate WM degeneration, as increased FA may also arise from non-pathological factors such as reduced fibre crossing or altered cellularity [Armstrong et al., 2024]. Consistent with this interpretation, several brain measures associated with CPcal in our study - including hippocampal, nucleus accumbens, and amygdala volumes, as well as mean diffusivity - were reported to mediate up to 42.9% of the association between CP volume and incident dementia in the same cohort [Yu et al., 2026]. These findings suggest that CPcal and CP volume may be associated with overlapping brain changes.

We examined maternal, paternal, and either-parent history of ADRD separately, as previous studies have reported differences in dementia-related imaging markers according to parental history [Mosconi et al., 2007; Mosconi et al., 2010]. In our study, maternal history was associated with greater CPcal in the lateral ventricle (d = 0.063), whereas paternal and either-parent history were not. This finding should be considered preliminary, given the small effect size and the potential for maternal history to be more reliably reported than paternal history in self-report cohorts. It also contrasts with prior studies reporting no difference in CPcal between Alzheimer’s disease [Friedland et al., 1990; Wu and Swaab, 2005]. However, the CP contributes to CSF homeostasis and amyloid-β clearance and has been associated with an Alzheimer’s disease subtype characterized by CP dysfunction [Tijms et al., 2024]. A recent longitudinal UK Biobank study linked greater CP volume to cognitive decline and incident dementia [Yu et al., 2026], supporting further investigation of CP-related markers in dementia risk despite the weak family-history association observed in our study.

Several limitations should be considered. First, this is a cross-sectional study. Although CPcal was higher in participants whose diagnoses were recorded after imaging, we cannot establish whether calcification precedes disease onset, and reverse causation cannot be excluded. Second, independent replication was not possible, to our knowledge, as no comparable large-scale dataset currently combines QSM with the phenotypic and diagnostic information available in the UK Biobank. Finally, the healthier profile of UK Biobank imaging participants and restriction of the family-history analysis to self-reported may limit generalizability. Future longitudinal studies should examine CPcal progression and its association with incident disorders.

In conclusion, CPcal is a common, quantifiable, and relatively understudied feature of the aging brain that can be assessed at scale using QSM without ionizing radiation. Our findings indicate that CPcal captures information distinct from CP volume, is associated with cardiometabolic and vascular burden across diagnostic categories, and shows its strongest associations in the third ventricle. This region remains comparatively underexplored in the CP imaging literature.

## Acknowledgments

This work was supported by the National Institutes of Health (R01AG087513, RF1NS136995, R01AG059874 and R01MH134004) Data were provided through the UK Biobank Resource (Application No. 11559). Computational analyses were funded partially through the support of NIH grant S10OD032285.

