## Supplementary_Material for "Regional choroid plexus calcifications and their associations with aging, brain structure, and disease"

### Supplementary Materials

#### ICD-10 disease classification

**Table S1 |** International Statistical Classification of Diseases (ICD-10), and corresponding level-1 that were analyzed in the study

| ICD-10 chapter (code range) | ICD-10 (Level 1) | |
|---|---|---|
| | Code range | Description |
| Chapter IV Endocrine, nutritional and metabolic diseases | ICD-10: E00-E07; SR: 1226, 1610, 1225, 1522, 1428, 1224 | Disorders of thyroid gland |
| | ICD-10: E10-E14; SR: 1222, 1223, 1220 | Diabetes mellitus |
| | ICD-10: E10; SR: 1222 | Insulin-dependent diabetes mellitus |
| | ICD-10: E11; SR: 1223 | Non-insulin-dependent diabetes mellitus |
| | ICD-10: E14; SR: 1220 | Unspecified diabetes mellitus |
| | ICD-10: E20-E35; SR: 1229, 1611, 1429, 1431, 1237, 1430, 1521, 1239, 1232, 1233, 1234, 1350, 1551, 1432 | Disorders of other endocrine glands |
| | ICD-10: E50-E64 | Other nutritional deficiencies |
| | ICD-10: E65-E68 | Obesity and other hyperalimentation |
| | ICD-10: E66 | Obesity |
| | ICD-10: E70-E90; SR: 1473, 1507, 1496 | Metabolic disorders |
| | ICD-10: E78; SR: 1473 | Disorders of lipoprotein metabolism and other lipidaemias |
| | ICD-10: E86 | Volume depletion |
| Chapter V Mental and behavioral disorders | ICD-10: F00-F09; SR: 1243 | Organic, including symptomatic, mental disorders |
| | ICD-10: F10-F19; SR: 1408, 1409 | Mental and behavioral disorders due to psychoactive substance use |
| | ICD-10: F17 | Mental and behavioural disorders due to use of tobacco |
| | ICD-10: F30-F39; SR: 1286 | Mood [affective] disorders |
| | ICD-10: F32; SR: 1286 | Depressive episode |
| | ICD-10: F40-F48; SR: 1469, 1614, 1615 | Neurotic, stress-related and somatoform disorders |
| | ICD-10: F41 | Other anxiety disorders |
| | ICD-10: F50-F54, F59; SR: 1470, 1531 | Behavioral syndromes associated with physiological disturbances and physical factors |
| Chapter VI Diseases of the nervous system | ICD-10: G00-G06, G08, G09; SR: 1247, 1246, 1245, 1248 | Inflammatory diseases of the central nervous system |
| | ICD-10: G20, G21, G23-G25; SR: 1262, 1525 | Extrapyramidal and movement disorders |
| | ICD-10: G35-G37; SR: 1261, 1397 | Demyelinating diseases of the central nervous system |
| | ICD-10: G40, G41, G43-G47; SR: 1264, 1265, 1082, 1123, 1616 | Episodic and paroxysmal disorders |
| | ICD-10: G43; SR: 1265 | Migraine |
| | ICD-10: G47; SR: 1123, 1616 | Sleep disorders |
| | ICD-10: G50-G59; SR: 1523, 1250, 1249, 1541 | Nerve, nerve root and plexus disorders |
| | G56 | Mononeuropathies of upper limb |

| | | |
|---|---|---|
| | ICD-10: G60-G64; SR: 1256, 1255, 1254 | Polyneuropathies and other disorders of the peripheral nervous system |
| | ICD-10: G90-G99; SR: 1251, 1434 | Other disorders of the nervous system |
| Chapter IX Diseases of the circulatory system | ICD-10: I05-I09 | Chronic rheumatic heart diseases |
| | ICD-10: I10-I13, I15; SR: 1072 | Hypertensive diseases |
| | ICD-10: I10; SR: 1072 | Essential (primary) hypertension |
| | ICD-10: I20-I25; SR: 1074, 1075 | Ischemic heart diseases |
| | I21 | Acute myocardial infarction |
| | I25 | Chronic ischaemic heart disease |
| | ICD-10: I26-I28; SR: 1093 | Pulmonary heart disease and diseases of pulmonary circulation |
| | I26 | Pulmonary embolism |
| | ICD-10: I30-I38, I40, I42-I51; SR: 1589, 1080, 1590, 1488, 1079, 1588, 1484, 1487, 1471, 1483, 1077, 1486, 1076, 1426 | Other forms of heart disease |
| | I48 | Atrial fibrillation and flutter |
| | I50 | Heart failure |
| | ICD-10: I60, I61, I62, I63, I64, I65, I66, I67, I68;<br>SR: 1086, 1583, 1081, 1425 | Cerebrovascular diseases |
| | ICD-10: I64; SR: 1081 | Stroke, not specified as haemorrhage or infarction |
| | ICD: I70-I74, I77-I79; SR: 1492, 1591, 1592, 1067, 1087, 1561, 1088 | Diseases of arteries, arterioles and capillaries |
| | ICD-10: I80-I89; SR: 1094, 1068, 1494, 1593, 1141, 1495 | Diseases of veins, lymphatic vessels and lymph nodes, not elsewhere classified |
| | I83 | Varicose veins of lower extremities |

The 10th revision of the International Statistical Classification of Diseases (ICD-10), Self-report (SR), Death record (DR)

#### Diseases progression: level-1 diagnostic and disorders associations

Within each ICD-10 chapter, associations between CPcal and individual-level 1 diagnoses and disorders for the progression analysis are reported below

##### Endocrine, nutritional and metabolic diseases

Participants with endocrine and metabolic disease (Table S2), diabetes mellitus were associated with greater CPcal in the 3rdV ($d = 0.3396$, $p = 1.82 \times 10^{-5}$); non-insulin-dependent diabetes mellitus was associated with greater CPcal in the 3rdV ($d = 0.2924$, $p = 6.97 \times 10^{-4}$); metabolic disorders was associated with greater CPcal in the 3rdV ($d = 0.1097$, $p = 0.0061$); disorders of lipoprotein metabolism was associated with greater CPcal in the 3rdV ($d = 0.1318$, $p = 0.0049$).

**Table S2 |** Associations between choroid plexus calcification (CPcal) volumes and level-1 diagnostic categories within ICD-10 chapter IV (Endocrine, nutritional and metabolic diseases). Significant level-1 categories are further broken down into specific ICD-10 disorders.

| ICD-10 chapters | CPcal Region | Effect Size | β | SE | 95% CI | P-value |
|---|---|---|---|---|---|---|
| Disorders of thyroid gland (N=406) | **LV** | 0.1396 | 0.1405 | 0.0711 | (9.39x10-4, 0.2800) | **0.0485** |

| | | | | | | |
|---|---|---|---|---|---|---|
| | **3rdV** | 0.1337 | 0.1436 | 0.0759 | (-0.0054, 0.2925) | 0.0589 |
| | **4thV*** | 1.2532 | 0.2257 | 0.1609 | (0.9147, 1.7196) | 0.1607 |
| Diabetes mellitus (N=329) | **LV** | 0.1674 | 0.1670 | 0.0784 | (0.0129, 0.3210) | **0.0337** |
| Diabetes mellitus (N=329) | **3rdV** | 0.3396 | 0.3434 | 0.0795 | (0.1872, 0.4995) | **1.82x10$^{-5}$** |
| Non-insulin-dependent diabetes mellitus (N=277) | **3rdV** | 0.2924 | 0.2906 | 0.0852 | (0.1232, 0.4579) | **6.97x10$^{-4}$** |
| Diabetes mellitus (N=329) | **4thV*** | 1.0980 | 0.0935 | 0.1796 | (0.7719, 1.5617) | 0.6027 |
| Other nutritional deficiencies (N=126) | **LV** | -0.0415 | -0.0455 | 0.1410 | (-0.3233, 0.2324) | 0.7475 |
| | **3rdV** | 0.0207 | 0.0219 | 0.1365 | (-0.2469, 0.2907) | 0.8726 |
| | **4thV*** | 0.7363 | -0.3061 | 0.3002 | (0.4072, 1.3251) | 0.3079 |
| Obesity and other hyperalimentation (N=418) | **LV** | 0.0971 | 0.0975 | 0.0699 | (-0.0397, 0.2347) | 0.1633 |
| | **3rdV** | 0.1397 | 0.1540 | 0.0767 | (0.0034, 0.3046) | **0.0450** |
| | **4thV*** | 1.1627 | 0.1508 | 0.1620 | (0.8467, 1.5984) | 0.3519 |
| Metabolic disorders (N=1258) | **LV** | 0.0704 | 0.0700 | 0.0397 | (-0.0079, 0.1479) | 0.0784 |
| Metabolic disorders (N=1258) | **3rdV** | 0.1097 | 0.1178 | 0.0429 | (0.0337, 0.2020) | **0.0061** |
| Disorders of lipoprotein metabolism and other lipidaemias (N=921) | **3rdV** | 0.1318 | 0.1422 | 0.0504 | (0.0433, 0.2412) | **0.0049** |
| Metabolic disorders (N=1258) | **4thV*** | 0.9163 | -0.0874 | 0.0904 | (0.7674, 1.0939) | 0.3338 |

**Uncorrected p ≤ critical FDR threshold *Uncorrected p ≤ 0.05

#### Mental and behavioral disorders

Participants diagnosed with mental and behavioral disorders (Table S3), disorders due to psychoactive substance use were associated with greater CPcal in the LV (d = 0.2489, p = 0.0097) and the 3rdV (d = 0.3853, p = 6.79x10$^{-5}$); tobacco use disorders were associated with greater CPcal in the LV (d = 0.2948, p = 0.0101) and the 3rdV (d = 0.4833, p = 2.95x10$^{-5}$). However, neurotic, stress-related and somatoform disorders and other anxiety disorders were not significantly associated with CPcal after FDR correction.

**Table S3 |** Associations between choroid plexus calcification (CPcal) volumes and level-1 diagnostic categories within ICD-10 chapter V (Mental and behavioral disorders). Significant level-1 categories are further broken down into specific ICD-10 disorders.

| **ICD-10 chapters** | **CPcal Region** | ***Effect Size*** | **β** | ***SE*** | ***95% CI*** | ***P-value*** |
|---|---|---|---|---|---|---|
| Mental and behavioural disorders due to psychoactive substance use (N=223) | **LV** | 0.2489 | 0.2646 | 0.1018 | (0.0645, 0.4647) | **0.0097** |
| Mental and behavioural disorders due to use of tobacco (N=159) | **LV** | 0.2948 | 0.3017 | 0.1166 | (0.0722, 0.5312) | **0.0101** |
| Mental and behavioural disorders due to psychoactive substance use (N=223) | **3rdV** | 0.3853 | 0.3923 | 0.0975 | (0.2006, 0.5839) | **6.79x10$^{-5}$** |
| Mental and behavioural disorders due to use of tobacco (N=159) | **3rdV** | 0.4833 | 0.4850 | 0.1144 | (0.2599, 0.7100) | **2.95x10$^{-5}$** |
| Mental and behavioural disorders due to psychoactive substance use (N=223) | **4thV*** | 1.1162 | 0.1099 | 0.2173 | (0.7290, 1.7109) | 0.6129 |
| Mood [affective] disorders (N=298) | **LV** | 0.0948 | 0.0982 | 0.0855 | (-0.0698, 0.2662) | 0.2515 |
| | **3rdV** | 0.0408 | 0.0461 | 0.0934 | (-0.1374, 0.2296) | 0.6218 |
| | **4thV*** | 1.0073 | 0.0073 | 0.1845 | (0.7016, 1.4472) | 0.9685 |
| Neurotic, stress-related and somatoform | **LV** | 0.1729 | 0.1924 | 0.0860 | (0.0235, 0.3612) | **0.0256** |

| | | | | | | |
|---|---|---|---|---|---|---|
| disorders (N=340) | 3rdV | 0.1352 | 0.1474 | 0.0843 | (-0.0180, 0.3128) | 0.0807 |
| Other anxiety disorders (N=307) | | 0.1559 | 0.1676 | 0.0875 | (-0.0042, 0.3394) | 0.0559 |
| Neurotic, stress-related and somatoform disorders (N=340) | 4thV* | 0.8609 | -0.1498 | 0.1716 | (0.6145, 1.2046) | 0.3827 |
| **Uncorrected p ≤ critical FDR threshold *Uncorrected p ≤ 0.05 | | | | | | |

#### Diseases of the nervous system

Participants diagnosed with nervous system disease, no disorders were significantly associated with CPcal after FDR correction; the episodic and paroxysmal disorders association reported previously did not reach significance (d = 0.1469, p = 0.0720) (Table S4).

**Table S4 |** Associations between choroid plexus calcification (CPcal) volumes and level-1 diagnostic categories within ICD-10 chapter VI (Diseases of the nervous system). Significant level-1 categories are further broken down into specific ICD-10 disorders.

| ICD-10 chapters | CPcal Region | *Effect Size* | β | *SE* | *95% CI* | *P-value* |
|---|---|---|---|---|---|---|
| Episodic and paroxysmal disorders (N=306) | LV | 0.0472 | 0.0498 | 0.0859 | (-0.1189, 0.2185) | 0.5627 |
| | 3rdV | 0.1469 | 0.1535 | 0.0852 | (-0.0138, 0.3208) | **0.0720** |
| | 4thV* | 1.2413 | 0.2162 | 0.1783 | (0.8756, 1.7624) | 0.2254 |
| Nerve, nerve root and plexus disorders (N=264) | LV | 0.0965 | 0.1117 | 0.1016 | (-0.0880, 0.3113) | 0.2724 |
| | 3rdV | 0.1204 | 0.1384 | 0.1010 | (-0.0600, 0.3369) | 0.1712 |
| | 4thV* | 1.0162 | 0.0161 | 0.1960 | (0.6917, 1.4927) | 0.9346 |
| **Uncorrected p ≤ critical FDR threshold *Uncorrected p ≤ 0.05 | | | | | | |

#### Diseases of the circulatory system

Participants diagnosed with circulatory system disease (Table S5), hypertensive diseases were associated with greater CPcal in the LV (d = 0.1134, p = 0.0026) and the 3rdV (d = 0.1677, p = $8.85 \times 10^{-6}$); essential (primary) hypertension in the 3rdV (d = 0.1655, p = $1.21 \times 10^{-5}$).

Pulmonary heart disease and diseases of pulmonary circulation were associated with greater CPcal in the 3rdV (d = 0.3806, p = 0.0039). Atrial fibrillation and flutter was no longer significantly associated with CPcal after FDR correction. No significant associations were observed for chronic rheumatic heart disease, ischaemic heart disease, diseases of arteries, or diseases of veins.

**Table S5 |** Associations between choroid plexus calcification (CPcal) volumes and level-1 diagnostic categories within ICD-10 chapter IX (Diseases of the circulatory system). Significant level-1 categories are further broken down into specific ICD-10 disorders.

| ICD-10 chapters | CPcal Region | *Effect Size* | β | *SE* | *95% CI* | *P-value* |
|---|---|---|---|---|---|---|
| Chronic rheumatic heart diseases (N=141) | LV | -0.1728 | -0.1786 | 0.1254 | (-0.4254, 0.0682) | 0.1554 |

| | | | | | | |
|---|---|---|---|---|---|---|
| | **3rdV** | -0.1466 | -0.1579 | 0.1306 | (-0.4150, 0.0992) | 0.2277 |
| | **4thV*** | 1.1339 | 0.1257 | 0.2786 | (0.6570, 1.9636) | 0.6519 |
| Hypertensive diseases (N=1415) | **LV** | 0.1134 | 0.1169 | 0.0388 | (0.0408, 0.1931) | **0.0026** |
| Essential (primary) hypertension (N=1409) | | 0.1097 | 0.1132 | 0.0390 | (0.0368, 0.1896) | **0.0037** |
| Hypertensive diseases (N=1415) | **3rdV** | 0.1677 | 0.1751 | 0.0393 | (0.0980, 0.2523) | **$8.85\times10^{-6}$** |
| Essential (primary) hypertension (N=1409) | | 0.1655 | 0.1729 | 0.0394 | (0.0956, 0.2502) | **$1.21\times10^{-5}$** |
| Hypertensive diseases (N=1415) | **4thV*** | 0.9719 | -0.0285 | 0.0851 | (0.8225, 1.1484) | 0.7379 |
| Ischaemic heart diseases (N=598) | **LV** | -0.0050 | -0.0051 | 0.0595 | (-0.1218, 0.1116) | 0.9316 |
| | **3rdV** | 0.0202 | 0.0209 | 0.0601 | (-0.0969, 0.1387) | 0.7279 |
| | **4thV*** | 0.9100 | -0.0943 | 0.1298 | (0.7054, 1.1737) | 0.4675 |
| Pulmonary heart disease and diseases of pulmonary circulation (N=123) | **LV** | 0.1284 | 0.1376 | 0.1398 | (-0.1379, 0.4131) | 0.3261 |
| | **3rdV** | 0.3806 | 0.4023 | 0.1379 | (0.1306, 0.6739) | **0.0039** |
| | **4thV*** | 2.0710 | 0.7280 | 0.2952 | (1.1676, 3.7262) | **0.0137** |
| Other forms of heart disease (N=1031) | **LV** | 0.0895 | 0.0914 | 0.0451 | (0.0030, 0.1798) | 0.0428 |
| | **3rdV** | 0.1019 | 0.1048 | 0.0454 | (0.0158, 0.1938) | **0.0211** |
| Atrial fibrillation and flutter (N=464) | | 0.1522 | 0.1526 | 0.0662 | (0.0227, 0.2825) | **0.0214** |
| Other forms of heart disease (N=1031) | **4thV*** | 0.9685 | -0.0320 | 0.0980 | (0.7992, 1.1736) | 0.7442 |
| Cerebrovascular diseases (N=231) | **LV** | 0.0937 | 0.0887 | 0.0890 | (-0.0862, 0.2636) | 0.3197 |
| | **3rdV** | 0.1896 | 0.1887 | 0.0936 | (0.0047, 0.3727) | **0.0444** |
| | **4thV*** | 1.2421 | 0.2168 | 0.2129 | (0.8185, 1.8878) | 0.3084 |
| Diseases of arteries, arterioles and capillaries (N=206) | **LV** | 0.1417 | 0.1392 | 0.0980 | (-0.0534, 0.3319) | 0.1561 |
| | **3rdV** | 0.0185 | 0.0188 | 0.1015 | (-0.1807, 0.2183) | 0.8533 |
| | **4thV*** | 1.1252 | 0.1180 | 0.2204 | (0.7304, 1.7349) | 0.5924 |
| Diseases of veins, lymphatic vessels and lymph nodes, not elsewhere classified (N=241) | **LV** | 0.0925 | 0.1027 | 0.1023 | (-0.0984, 0.3038) | 0.3162 |
| | **3rdV** | -0.0290 | -0.0305 | 0.0971 | (-0.2213, 0.1602) | 0.7534 |
| | **4thV*** | 1.3015 | 0.2635 | 0.2087 | (0.8646, 1.9617) | 0.2068 |

**Uncorrected p ≤ critical FDR threshold *Uncorrected p ≤ 0.05

#### Quantile regression and sensitivity analyses

We also used quantile regression for the lateral- and third-ventricle outcomes. Age and cardiovascular risk factors were associated with calcification across the distribution, but the strength of these associations varied by quantile (Figure S1).

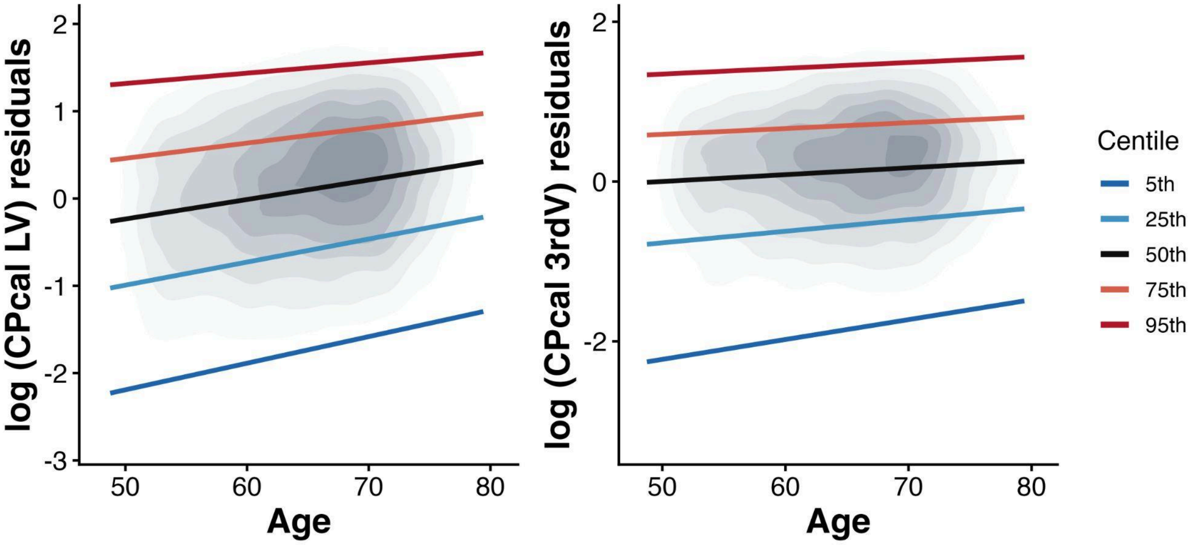


**Figure S1** Quantile regression centile curves for choroid plexus calcification in the lateral (left) and third (right) ventricles. Colored lines represent the fitted 5th, 25th, 50th, 75th and 95th centiles of covariate-adjusted CPcal as a function of age, overlaid on the density of data.

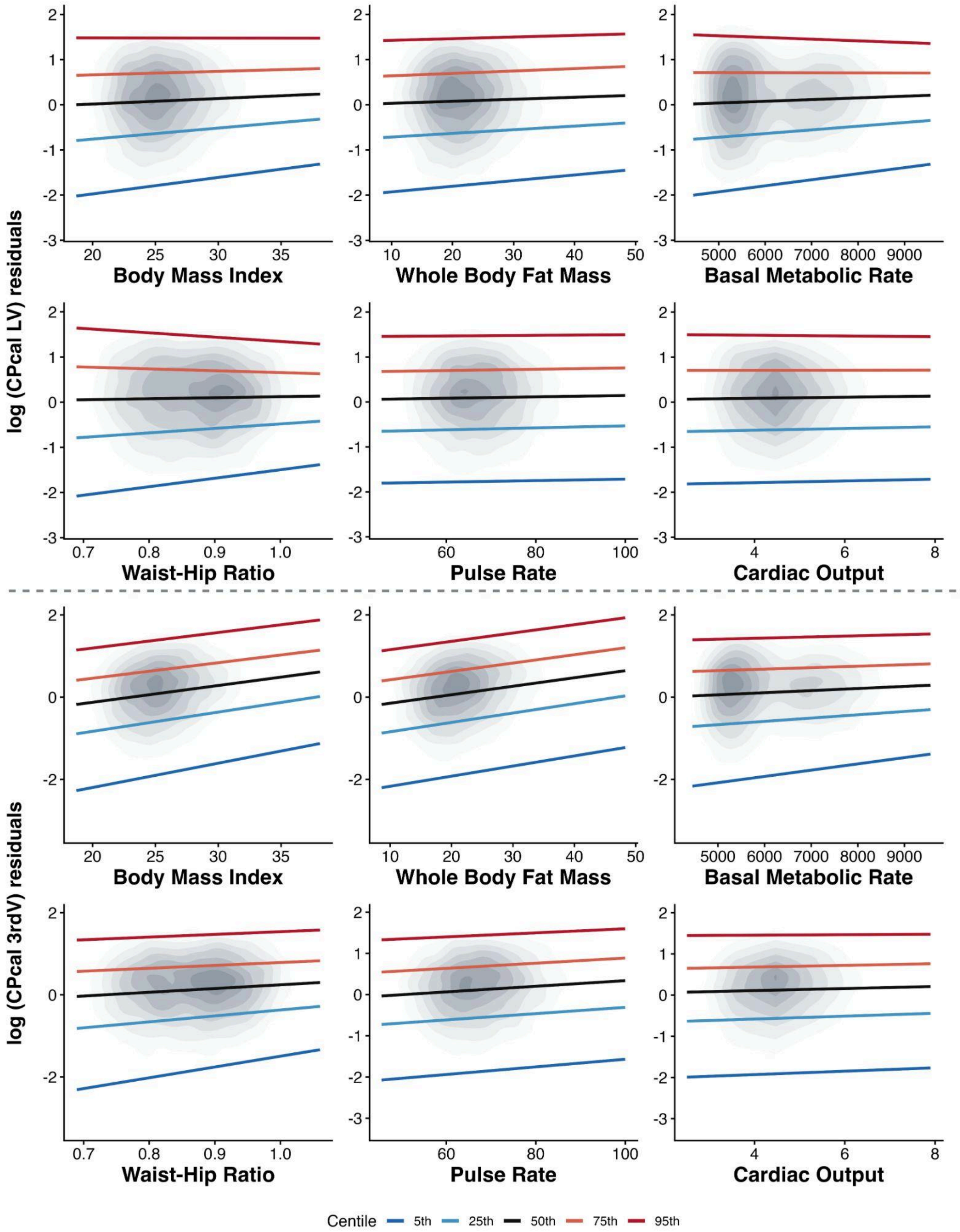
log (CPcal LV) residuals
Body Mass Index
Whole Body Fat Mass
Basal Metabolic Rate
Waist-Hip Ratio
Pulse Rate
Cardiac Output
log (CPcal 3rdV) residuals
Body Mass Index
Whole Body Fat Mass
Basal Metabolic Rate
Waist-Hip Ratio
Pulse Rate
Cardiac Output
Centile
5th
25th
50th
75th
95th

**Figure S2** Quantile regression centile curves for choroid plexus calcification in the lateral (left) and third (right) ventricles. Colored lines represent the fitted 5th, 25th, 50th, 75th and 95th centiles of covariate-adjusted CPcal as a function of cardiovascular risk factors, overlaid on the density of data. The risk factors body mass index, whole body fat mass, basal metabolic rate, waist hip ratio, pulse rate and cardiac output.

The third ventricle CPcal showed the strongest and most consistent disease associations. To determine whether these associations were related to ventricular enlargement, we adjusted each model for third ventricle volume. Of the 29 significant associations, 27 (93%) remained significant (Figure 6). These findings suggest that third ventricle CPcal is largely independent of ventricular volume and unlikely to reflect partial volume artifacts.

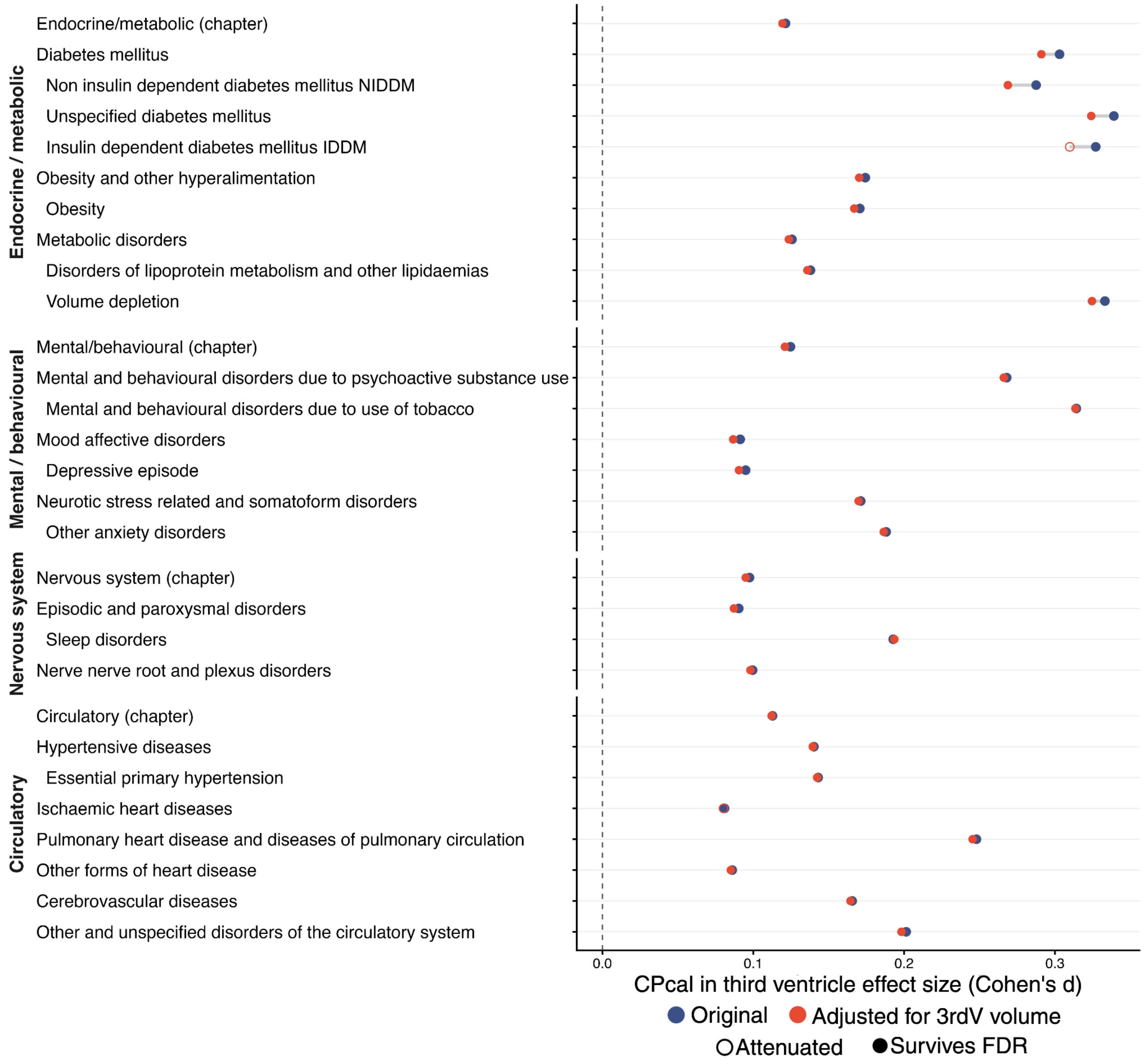


**Figure S3** Third ventricle sensitivity analysis. Effect sizes for diseases originally associated with third ventricle choroid plexus calcification are shown before (navy) and after (red) adjustment for third ventricle volume. Open circles indicate associations that became non-significant after adjustment. Disorders are grouped by ICD-10 chapter and ordered as in the main paper.

### Disease and Risk Factor Associations with CPcal

#### Quantile regression results: effect size

**Table S06 |** Quantile regression effect sizes for the association between choroid plexus calcification (CPcal) and ICD-10 diagnostic chapters in the lateral (LV) and third (3rdV) ventricles across calcification centiles. Bold values with an asterisk (**) indicate significance after Benjamini-Hochberg FDR correction (q < 0.05); * denotes uncorrected p < 0.05.

| ICD-10 chapters | CPcal Region | 5th | 25th | 50th | 75th | 95th |
|---|---|---|---|---|---|---|
| Endocrine, nutritional and metabolic diseases (N=7939) | LV | 0.0590 | 0.0361 | 0.0372 | 0.0294 | 0.0400 |
| | 3rdV | 0.1160$^{*}$ | **0.1429**$^{**}$ | **0.1087**$^{**}$ | **0.1111**$^{**}$ | **0.1431**$^{**}$ |
| Mental and behavioral disorders (N=4589) | LV | 0.1141 | **0.0741**$^{**}$ | **0.0605**$^{**}$ | 0.0254 | 0.0429 |
| | 3rdV | **0.2483**$^{**}$ | **0.1300**$^{**}$ | **0.0879**$^{**}$ | **0.1380**$^{**}$ | **0.1251**$^{**}$ |
| Diseases of the nervous system (N=5086) | LV | 0.0594 | 0.0536 | **0.0873**$^{**}$ | 0.0416 | 0.0506 |
| | 3rdV | 0.1084$^{*}$ | **0.1250**$^{**}$ | **0.1014**$^{**}$ | **0.0845**$^{**}$ | **0.0988**$^{**}$ |
| Diseases of the circulatory system (N=8248) | LV | 0.0390 | 0.0402 | 0.0234 | 0.0268 | 0.0333 |
| | 3rdV | **0.1518**$^{**}$ | **0.1281**$^{**}$ | **0.0972**$^{**}$ | **0.1039**$^{**}$ | **0.0970**$^{**}$ |

#### Endocrine, nutritional and metabolic diseases

**Table S07 |** Quantile regression effect sizes for the association between choroid plexus calcification (CPcal) and endocrine, nutritional, and metabolic diagnoses in the lateral (LV) and third (3rdV) ventricles across calcification centiles. Bold values with an asterisk (**) indicate significance after Benjamini-Hochberg FDR correction (q < 0.05); * denotes uncorrected p < 0.05.

| ICD-10 chapters | CPcal Region | 5th | 25th | 50th | 75th | 95th |
|---|---|---|---|---|---|---|
| Disorders of thyroid gland (N=2350) | LV | -0.0022 | 0.0105 | 0.0145 | 0.0151 | 0.0644 |
| | 3rdV | 0.0505 | 0.0610 | 0.0317 | 0.0368 | 0.0706 |
| Diabetes mellitus (N=1366) | LV | 0.1025 | **0.1462**$^{**}$ | **0.1248**$^{**}$ | 0.0647 | -0.0444 |
| Non-insulin-dependent diabetes mellitus (N=1055) | | 0.1631 | 0.1244$^{*}$ | 0.1302$^{*}$ | 0.0561 | -0.0508 |
| Unspecified diabetes mellitus (N=913) | | 0.1504 | 0.1639$^{*}$ | **0.1509**$^{**}$ | 0.0582 | 0.0012 |
| Diabetes mellitus (N=1366) | 3rdV | **0.3469**$^{**}$ | **0.3567**$^{**}$ | **0.2563**$^{**}$ | **0.2270**$^{**}$ | **0.2412**$^{**}$ |
| Insulin-dependent diabetes mellitus (N=130) | | **0.7812**$^{**}$ | 0.4337$^{*}$ | 0.1158 | 0.1128 | **0.3095**$^{**}$ |
| Non-insulin-dependent diabetes mellitus (N=1055) | | **0.3701**$^{**}$ | **0.3686**$^{**}$ | **0.2420**$^{**}$ | **0.2147**$^{**}$ | **0.2345**$^{**}$ |

| ICD-10 chapters | CPcal Region | 5th | 25th | 50th | 75th | 95th |
|---|---|---|---|---|---|---|
| Unspecified diabetes mellitus (N=913) | | **0.4194**** | **0.4077**** | **0.2611**** | **0.2231**** | **0.2438**** |
| Disorders of other endocrine glands (N=336) | LV | 0.1683 | 0.0339 | 0.0333 | 0.0276 | -0.0913 |
| | 3rdV | 0.5238 | 0.0017 | 0.1200 | 0.1595* | 0.2307 |
| Other nutritional deficiencies (N=222) | LV | 0.2481 | -0.0256 | 0.0136 | 0.0352 | -0.0651 |
| | 3rdV | -0.0669 | -0.0039 | -0.1203 | -0.0141 | -0.0686 |
| Obesity and other hyperalimentation (N=870) | LV | **0.3554**** | 0.1485* | 0.1172 | 0.0283 | -0.0010 |
| Obesity (N=867) | | **0.3554**** | 0.1361 | 0.1039 | 0.0314 | 0.0003 |
| Obesity and other hyperalimentation (N=870) | 3rdV | -0.1324 | **0.2079**** | **0.1555**** | **0.2398**** | **0.2483**** |
| Obesity (N=867) | | -0.1378 | **0.2100**** | **0.1481**** | **0.2362**** | **0.2483**** |
| Metabolic disorders (N=5840) | LV | 0.0729 | 0.0339 | 0.0312 | 0.0314 | 0.0622 |
| Disorders of lipoprotein metabolism and other lipidaemias (N=5435) | | 0.0992* | 0.0406 | 0.0558* | 0.0491* | 0.0402 |
| Metabolic disorders (N=5840) | 3rdV | 0.1237* | **0.1511**** | **0.1344**** | **0.1183**** | **0.1247**** |
| Disorders of lipoprotein metabolism and other lipidaemias (N=5435) | | **0.1670**** | **0.1627**** | **0.1416**** | **0.1159**** | **0.1228**** |
| Volume depletion (N=168) | | **1.1235**** | 0.2401 | 0.2578* | 0.2926* | 0.0447 |

#### Mental and behavioral disorders

**Table S08 |** Quantile regression effect sizes for the association between choroid plexus calcification (CPcal) and mental and behavioral diagnoses in the lateral (LV) and third (3rdV) ventricles across calcification centiles. Bold values with an asterisk (**) indicate significance after Benjamini-Hochberg FDR correction ($q < 0.05$); * denotes uncorrected $p < 0.05$.

| ICD-10 chapters | CPcal Region | 5th | 25th | 50th | 75th | 95th |
|---|---|---|---|---|---|---|
| Organic, including symptomatic, mental disorders (N=119) | LV | 0.2106 | **0.2518**** | 0.0402 | 0.3278 | 0.2643 |
| | 3rdV | 0.2660 | 0.3495 | 0.1152 | 0.2426 | 0.1196 |
| Mental and behavioral disorders due to psychoactive substance use (N=760) | LV | 0.0514 | 0.1537 | **0.1728**** | **0.1381**** | **0.1736**** |
| Mental and behavioral disorders due to use of tobacco (N=538) | | 0.2118 | 0.2058* | **0.2005**** | 0.1463 | 0.2018** |
| Mental and behavioral disorders due to psychoactive substance use (N=760) | 3rdV | 0.3724* | **0.2338**** | **0.2531**** | **0.3038**** | **0.2102**** |
| Mental and behavioral disorders due to use of tobacco (N=538) | | 0.2733 | **0.2638**** | **0.2574**** | **0.3030**** | **0.2460**** |
| Mood [affective] disorders (N=3138) | LV | 0.1372 | **0.1133**** | **0.1151**** | **0.0716**** | 0.0231 |
| Depressive episode (N=3112) | | 0.1371 | **0.1159**** | **0.1125**** | 0.0685* | 0.0296 |
| Mood [affective] disorders (N=3138) | 3rdV | 0.2018* | **0.1057**** | 0.0662 | **0.1142**** | 0.0799* |
| Depressive episode (N=3112) | | 0.2127* | **0.0993**** | **0.0773**** | **0.1154**** | 0.0841* |
| Neurotic, stress-related and somatoform disorders (N=1190) | LV | 0.2420* | **0.2018**** | **0.1587**** | 0.0998* | 0.0825 |
| Other anxiety disorders (N=681) | | -0.0174 | **0.2008**** | **0.1851**** | **0.1871**** | 0.0788 |

| | | | | | | |
|---|---|---|---|---|---|---|
| Neurotic, stress-related and somatoform disorders (N=1190) | 3rdV | 0.2525* | **0.1450**** | **0.1191**** | **0.2027**** | **0.1805**** |
| Other anxiety disorders (N=681) | | 0.3132* | 0.1930* | 0.0856 | **0.1863**** | 0.1057 |
| Behavioral syndromes associated with physiological disturbances and physical factors (N=226) | LV | -0.1424 | -0.0449 | **0.3158**** | **0.2298**** | -0.1748 |
| | 3rdV | 0.0906 | 0.1244 | 0.2313 | **0.1973**** | 0.1651 |

#### Diseases of the nervous system

**Table S09 |** Quantile regression effect sizes for the association between choroid plexus calcification (CPcal) and nervous system diagnoses in the lateral (LV) and third (3rdV) ventricles across calcification centiles. Bold values with an asterisk (**) indicate significance after Benjamini-Hochberg FDR correction (q < 0.05); * denotes uncorrected p < 0.05.

| ICD-10 chapters | CPcal Region | 5th | 25th | 50th | 75th | 95th |
|---|---|---|---|---|---|---|
| Inflammatory diseases of the central nervous system (N=201) | LV | 0.0468 | -0.1457 | -0.1735 | -0.1315 | -0.2106 |
| | 3rdV | -0.0145 | -0.1397 | -0.0391 | -0.0666 | **-0.2038**** |
| Extrapyramidal and movement disorders (N=174) | LV | -0.0499 | -0.0531 | 0.0018 | -0.0754 | -0.1720 |
| | 3rdV | 0.2948 | 0.0880 | 0.0003 | 0.0835 | 0.1944 |
| Demyelinating diseases of the central nervous system (N=130) | LV | **0.7403**** | 0.0314 | -0.0621 | -0.0487 | -0.1382 |
| | 3rdV | 0.1323 | 0.2742 | 0.3150 | **0.2412**** | -0.0752 |
| Episodic and paroxysmal disorders (N=3060) | LV | -0.0233 | 0.0584 | **0.0814**** | 0.0595* | 0.0523 |
| | 3rdV | 0.0398 | 0.1172* | **0.0971**** | **0.0849**** | **0.1052**** |
| Migraine (N=2125) | | 0.0979 | 0.1124* | 0.0886* | 0.0875* | 0.1046* |
| Sleep disorders (N=443) | | 0.0869 | 0.3059* | **0.2887**** | 0.1720* | 0.1155 |
| Nerve, nerve root and plexus disorders (N=1440) | LV | 0.1161 | 0.0930 | **0.1349**** | 0.0855 | **0.1352**** |
| Mononeuropathies of upper limb (N=794) | | 0.2493 | 0.1304 | 0.1544* | **0.1771**** | **0.2876**** |
| Other mononeuropathies (N=951) | | 0.1971 | 0.1296 | 0.1452* | **0.1614**** | **0.2497**** |
| Nerve, nerve root and plexus disorders (N=1440) | 3rdV | 0.2296* | 0.1558* | 0.0709 | 0.0811 | 0.0486 |
| Mononeuropathies of upper limb (N=794) | | 0.1053 | 0.0445 | 0.1225* | **0.1437**** | 0.0247* |
| Other mononeuropathies (N=951) | | 0.1592 | 0.1080 | 0.1383 | 0.1014* | 0.0201* |
| Polyneuropathies and other disorders of the peripheral nervous system (N=247) | LV | **0.3813**** | 0.2485* | 0.0903 | 0.0278 | 0.0786 |
| | 3rdV | **-0.4852**** | 0.1284 | 0.1045 | -0.0301 | -0.0706 |
| Other disorders of the nervous system (N=317) | LV | 0.2904 | 0.1660 | 0.1523 | **0.2782**** | 0.1320 |
| | 3rdV | -0.0285 | 0.0974 | **0.1846**** | 0.1554 | 0.1392 |

#### Diseases of the circulatory system

**Table S10 |** Quantile regression effect sizes for the association between choroid plexus calcification (CPcal) and circulatory system diagnoses in the lateral (LV) and third (3rdV) ventricles across calcification centiles. Bold values with an asterisk (**) indicate significance after Benjamini-Hochberg FDR correction (q < 0.05); * denotes uncorrected p < 0.05.

| ICD-10 chapters | CPcal Region | 5th | 25th | 50th | 75th | 95th |
|---|---|---|---|---|---|---|
| Chronic rheumatic heart diseases (N=232) | LV | -0.0487 | 0.0342 | -0.0699 | -0.0056 | -0.1981 |
| | 3rdV | -0.1070 | 0.0556 | 0.0565 | 0.0393 | 0.1553 |
| Hypertensive diseases (N=5055) | LV | 0.0609 | 0.0618* | 0.0549* | 0.0541* | 0.0518 |
| Essential (primary) hypertension (N=5052) | | 0.0619 | 0.0619* | 0.0541* | 0.0557* | 0.0530 |
| Hypertensive diseases (N=5055) | 3rdV | 0.1241 | **0.1705**** | **0.1281**** | **0.1298**** | **0.1262**** |
| Essential (primary) hypertension (N=5052) | | 0.1244* | **0.1745**** | **0.1330**** | **0.1337**** | **0.1237**** |
| Ischaemic heart diseases (N=1913) | LV | 0.0983 | 0.0506 | 0.0315 | -0.0127 | 0.0340 |
| | 3rdV | 0.0343 | 0.0592 | 0.0798* | 0.0904* | **0.1492**** |
| Chronic ischaemic heart disease (N=1447) | | 0.0978 | 0.0690 | 0.1006* | 0.1103* | 0.0711 |
| Pulmonary heart disease and diseases of pulmonary circulation (N=359) | LV | 0.0737 | 0.2085 | 0.1203 | 0.0952 | 0.1531 |
| | 3rdV | 0.3752 | 0.3226* | **0.2361**** | 0.2508 | 0.0597 |
| Pulmonary embolism (N=334) | | 0.4072 | 0.3224* | 0.2491* | 0.3101* | 0.1370 |
| Other forms of heart disease (N=2384) | LV | -0.0123 | 0.0740 | 0.0269 | -0.0073 | -0.0203 |
| | 3rdV | 0.1269 | 0.0856* | 0.0604 | **0.1065**** | **0.1124**** |
| Atrial fibrillation and flutter (N=1118) | | 0.1369 | 0.1730* | 0.0714 | 0.0878* | 0.1211 |
| Heart failure (N=286) | | 0.0542 | 0.2547* | 0.1769* | 0.2387* | 0.0754 |
| Cerebrovascular diseases (N=636) | LV | -0.0253 | 0.0831 | 0.1526* | 0.0820 | 0.1897* |
| | 3rdV | 0.0726 | **0.2467**** | **0.1646**** | 0.0937 | 0.0800 |
| Stroke, not specified as haemorrhage or infarction (N=297) | | 0.0547 | **0.4350**** | 0.1824 | 0.1548 | -0.0951 |
| Diseases of arteries, arterioles and capillaries (N=585) | LV | -0.3072* | -0.1674* | 0.0464 | -0.0248 | 0.0800 |
| | 3rdV | 0.0966 | 0.1979* | 0.0909 | 0.0587 | 0.0439 |
| Diseases of veins, lymphatic vessels and lymph nodes, not elsewhere classified (N=3350) | LV | 0.0814 | 0.0386 | 0.0548 | 0.0159 | 0.0285 |
| | 3rdV | 0.0784 | 0.0850* | 0.0523 | **0.0613**** | -0.0005 |
| Varicose veins of lower extremities (N=1185) | | **0.2392**** | 0.1063 | 0.1314 | 0.1202 | 0.0241 |
